# Replication of SARS-CoV-2 Omicron lineages is defined by TMPRSS2 use in environments where ACE2 is complexed with solute carriers SLC6A19 and SLC6A20

**DOI:** 10.1101/2025.07.07.663433

**Authors:** Anupriya Aggarwal, Samantha Ognenovska, Timothy Ison, Christina Fichter, Vanessa Milogiannakis, Madeeha Afzal, Alberto Ospina Stella, Shafagh Waters, Camille Esneau, Nathan Bartlett, Stefan Pöhlmann, Markus Hoffmann, Louise M Burrell, Sheila K Patel, Sally Ellis, Michael Wehrhahn, Elena Martinez, Andrew Ginn, Melissa Churchill, Thomas Angelovich, William Rawlinson, Malinna Yeang, Jen Kok, Vitali Sintchenkov, Rhys Parry, Julian D Sng, Greg Neely, Cesar Moreno, Lipin Loo, Anthony D Kelleher, Fabienne Brilot, Alexander Khromykh, Stuart G Turville

**Author notes:** Equal author contribution (Anupriya Aggarwal, Samantha Ognenovska, Timothy Ison).

## Abstract

The Omicron variant of SARS-CoV-2 emerged in late 2021 and since then Omicron subvariants have continued to evolve and dominate globally. The viral S protein evolved towards highly efficient antibody evasion and replicative capacity in the upper respiratory tract resulting in high transmissibility. At the same time, the mutations acquired in the S protein diminish infection of the lung epithelium and pathogenic potential. The changing entry requirements for Omicron sub-lineages that lead to this shift in tropism remain poorly understood. We resolve the changing *replication* requirements of SARS-CoV-2 to be related to two distinct pools of ACE2. The first pool relates to ACE2’s role in the renin angiotensin system (RAS) and this pool can complex with TMPRSS2 (RAS-ACE2). The second pool relates to ACE2’s role as a protein solute carrier chaperone than cannot complex with TMPRSS2 (Chaperone ACE2). Here, we demonstrate that pre-Omicron lineages replicate in a TMPRSS2 dependent manner across both ACE2 pools, whilst Omicron lineages can only *spread and replicate using* chaperone ACE2. This provides a mechanistic basis for the evolving *infectivity* requirements of SARS-CoV-2 and furthermore provides approaches to track and monitor ACE2 utilizing coronaviruses.

**Graphical Abstract:** Mechanistic basis for shift in SARS-CoV-2 tropism with the arrival of Omicron. **A.** Chaperone ACE2 is defined structurally as a heterodimer of dimers with a solute carrier protein-SLC6A19 or SLC6A20. Here this ACE2 structure can exist uncomplexed from TMPRSS2 and enables TMPRSS2 use by both pre-Omicron and Omicron lineages. **B.** Renin Angiotensin ACE2 is defined by ACE2 with an exposed collectrin-like domain (CLD), which enables binding of TMPRSS2 or ADAM-17. Here ACE2 can form a complex with TMPRSS2 in a manner that allows pre-Omicron but not Omicron lineages to utilize TMPRSS2 to facilitate infection. Here Omicron lineages are heavily attenuated as they cannot use TMPRSS2 to spread. **C. to E.** Based on single cell profiles, ACE2 can exist as a chaperone with SLC6A20 in **C.** the Nasal Cavity or **D.** primarily as RAS-ACE2 in the lung to respond to acute lung injury. **E.** The largest pool of ACE2 in our body resides within the small intestine on enterocytes and this further facilitates replication in this tissue by pre-Omicron and Omicron lineages.

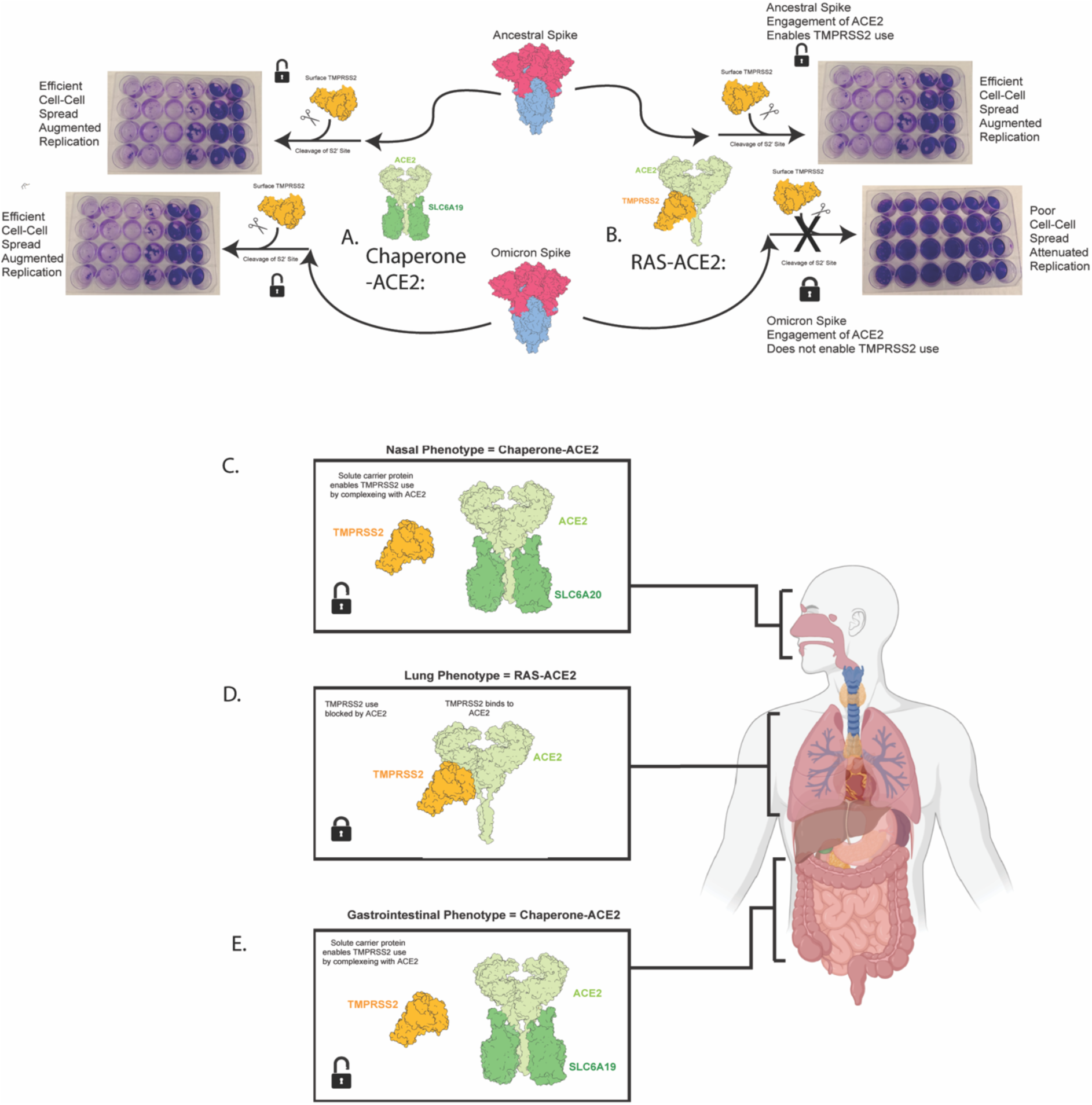

## Introduction

The four endemic coronaviruses 229E, OC43, NL63, and HKU1 circulate globally and usually cause mild to moderate upper respiratory tract infections. In contrast, severe acute respiratory syndrome coronavirus (SARS-CoV-1), Middle East respiratory syndrome (MERS-CoV) and SARS-CoV-2 can descend into the lower respiratory tract and cause severe respiratory disease. The manifestations of clinical disease are complex and influenced by prior vaccine and convalescent immunity and a greater understanding of the coronavirus cell and organ tropism is required to resolve the viral pathogenesis. The coronavirus spike protein (S) facilitates viral entry into target cells by binding to cellular receptors and fusing the viral membrane with a host cell membrane upon S protein cleavage-activation (priming) by a cellular protease. As a consequence, expression of receptor and protease are major determinants of coronavirus cell, organ and species tropism.

It was shown early in the COVID-19 pandemic that SARS-CoV-2, the causative agent of COVID-19, uses the cellular protein ACE2 as a primary receptor and the cellular serine proteases furin and TMPRSS2 for S protein activation. Activation was found to entail S protein cleavage at the furin cleavage site (FCS) within the S1/S2 cleavage site in infected cells during S protein passage through the constitutive secretory pathway, while TMPRSS2 cleaves the S protein at the S2’ site during viral entry (referred to as cleavage *in trans*), resulting in S protein activation for membrane fusion. Notably, TMPRSS2 cleavage at the S2’ site is dependent on previous furin-mediated cleavage of the FCS at the S1/S2 site ^1–3^. As auxiliary pathway, the SARS-CoV-2 S protein can also be activated upon cleavage by the cellular cysteine protease cathepsin L upon trafficking of virions into endo/lysosomes. However, the TMPRSS2 but not the cathepsin L-dependent entry pathway is operative in many primary cell models of the respiratory tract and TMPRSS2 was found to be essential for viral spread and pathogenesis in rodent models ^4–6^.

The emergence of SARS-CoV-2 variants of concern (VOC), Alpha, Beta, Gamma and Delta, was driven by mutations in the S protein that provided a fitness gain (Alpha) and augmented antibody evasion (Beta, Gamma and Delta). Fitness mutations like N501Y ^7,8^ increased affinity to ACE2, while mutations like P681R increased S protein cleavage at the S1/2 site and thus augmented lung cell entry^9–13^. The rapid replacement of the Delta variant by the Omicron variant in late 2021 constituted a major shift in SARS-CoV-2 evolution. The Omicron variant harbored an unusually high number of mutations in the S protein that allowed for an unprecedented level of antibody evasion. In parallel, the mutations shifted the viral tropism from the lower to the upper respiratory tract, allowing for high levels of transmission ^12–17^. However, the mutations also reduced TMPRSS2 usage in cell culture and diminished the ability to infect lung cells and to cause severe disease ^14,15,17–21^. On the other hand, studies with knock-out animals demonstrate TMPRSS2-dependence of infection by various Omicron subvariants, although this dependence is less pronounced than that observed for pre-Omicron variants ^5^. At present, the determinants controlling TMPRSS2 usage by Omicron subvariants and their impact on viral spread and pathogenesis are poorly understood.

Using clinical viral isolates that span the entire pandemic and subsequent endemic phase, we observe an evolution of entry requirements of Omicron subvariants. This evolution relates to the usage of two physiologically distinct pools of ACE2. ACE2 in the first pool can form a complex with TMPRSS2 and a related protease, ADAM-17, via its collectrin-like domain (CLD) as part of the renin angiotensin system (RAS, subsequently referred to RAS-ACE2).

The second ACE2 pool is generated by the formation of a dimer of heterodimers of ACE2 with one of two solute carriers, SLC6A19 and SLC6A20, hereon referred to as Chaperone ACE2. As the two solute carriers are complexed with the ACE2 CLD, TMPRSS2 can no longer form a complex with ACE2. While infection of pre-Omicron variants can be augmented by the presence of both ACE2 pools, replication of Omicron subvariants is attenuated in the presence of ACE2 pools that can form complexes with TMPRSS2. In this latter setting attenuation is mediated by the poor use of TMPRSS2 in this complex during cell-cell spread.

## Results

### High expression of ACE2 in TMPRSS2-positive cells augments infection by pre-Omicron variants but attenuates infection by Omicron lineages

The SARS-CoV-2 variant circulating at the beginning of the COVID-19 pandemic in late 2019 (hereon referred to as the Ancestral Clade A) and subsequent variants of concern (VOC) Alpha to Delta can employ TMPRSS2 with high efficiency for cell entry if co-expressed with ACE2 ^22,23^. In contrast, Omicron sub-lineages showed reduced capacity to employ TMPRSS2 for cell entry, at least in various *in vitro* settings ^19–21,24,25^. The latter has reduced the ability to isolate, propagate and work with primary clinical isolates for continued surveillance and clinical efforts focused on next generation vaccines and therapeutics. In order to identify cell systems that allow for robust replication of both pre- and post-Omicron variants (Fig. 1A), we screened several cell lines with various levels of ACE2 and TMPRSS2 expression (Table I) for permissiveness to SARS-CoV-2 infection. For this, we employed a previously described assay ^17,26,27^, which measures the reduction of the number of cell nuclei in infected cultures due to virus induced cytopathic effects. Although all cell lines were susceptible to infection by a broad spectrum of pre- and post-Omicron variants, the cell line that was most efficiently and comparably infected by all viruses studied were VeroE6-TMPRSS2^high^ cells (Fig. S1. A-G; V-TMPRSS2(2)). Furthermore, use of the TMPRSS2 inhibitor Nafamostat showed that infection by all variants was dependent on TMPRSS2 (Fig. 1B-D). Notably, increasing ACE2 expression in VeroE6-^TMPRSS2hi^ cells by stable lentiviral transduction, to produce the V-ACE2^high^-TMPRSS2^high^ cell line, augmented infection by all pre-Omicron variants but reduced infection by all Omicron sub-lineages (Fig. 1F-I). Interestingly, this attenuation was more pronounced in Omicron sub-lineages circulating in 2023 compared to earlier sub-lineages circulating in 2022 (Fig. 1F-I). These results suggested that high levels of ACE2 expression in a TMPRSS2-positive cellular background can significantly reduce infection by Omicron subvariants, which may reflect a progressive change in their ability to co-utilize ACE2 and TMPRSS2 to mediate infection. In addition, attenuation groups major lineages of Omicrons each year with progressive steps in attenuation in 2022 viral lineages (moderate attenuation; < 5 fold drop in titer), 2023 lineages (high attenuation; > 5 fold drop in titer) and 2024 lineages (very high attenuation; > 10 fold drop in titer).

**Table I.** Cell lines initially screened for SARS-CoV-2 isolation across pre-Omicron and Omicron lineages.

| Cell Line | Parent line | TMPRSS2 | ACE2 | Source |
| --- | --- | --- | --- | --- |
| HAT-24 | HEK | low | high | <sup>17</sup> |
| Calu-3 | - | low* | low* | ATCC HTB-55 |
| VeroE6 | Vero | - | low* | ATCC CRL-1586 |
| VeroE6-TMPRSS2 | VeroE6 | moderate | low* | <sup>3</sup> |
| VeroE6-TMPRSS2<sup>high</sup> | VeroE6 | high | low* | <sup>28</sup> |
| V-ACE2<sup>high</sup>-TMPRSS2<sup>high</sup> | VeroE6-TMPRSS2<sup>high</sup> | high | high | this study |

**Figure 1.**
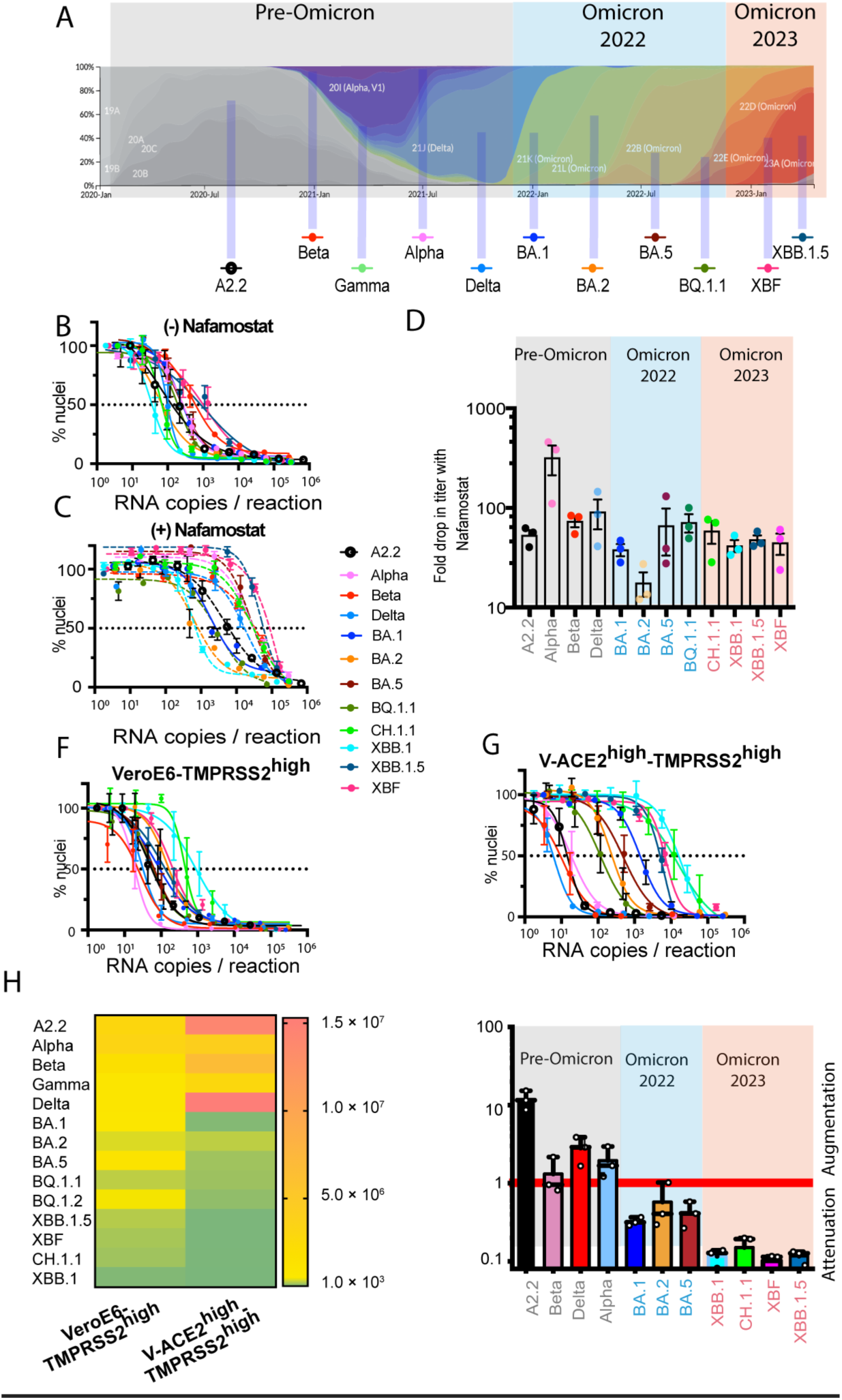
ACE2 and TMPRSS2 co-expression attenuates Omicron lineage use of TMPRSS2. **A.** Frequency of SARS-CoV-2 variants (nextstrain.org/sars-cov-2). Pre-Omicron variants are indicated in grey, while Omicron subvariants circulating during 2022 and 2023 are indicated in blue and pink, respectively. **B-C.** Titrations of 12 primary SARS-CoV-2 isolates in the VeroE6-TMPRSS2^high^ cell line in the presence of vehicle control (dimethyl sulfoxide; DMSO, panel B) or the TMPRSS2 inhibitor Nafamostat (20 µM, panel C). **D.** Summary of fold reduction in titers in the presence of Nafamostat from B-C; each data point represents an independent experiment. Each variant period is shaded in grey, blue and pink to align with variant periods outlined in A. **E.** Surface expression of ACE2 in the V-ACE2^high^TMPRSS2^high^ cell line, measured using flow cytometry, relative to other cell lines presented in Figure S1 B-H. **F-G.** Viral titers of 12 primary SARS-CoV-2 isolates in the F. VeroE6-TMPRSS2^high^ cell line and G. V-ACE2^high^TMPRSS2^high^ cell line. Left and right shifts in the sigmoidal curves indicate attenuation and augmentation of viral entry, respectively. **H.** Summary of viral titers from F-G. **I.** Replication of variants in the V-ACE2^high^TMPRSS2^high^ was compared relative to the parental VeroE6-TMPRSS2^high^ (2). (V-ACE2^high^TMPRSS2^high^)/ VeroE6-TMPRSS2^high^ titer) establishes levels of augmentation or attenuation across variant eras outlined in A. The red line is where titers are equivalent in each cell.

To investigate whether the observed attenuation of Omicron sub-lineages upon ACE2 expression was related to TMPRSS2, we also expressed high levels of ACE2 in a TMPRSS2-negative cellular background. For this, we employed the Affinofile system^29^, which allowed for inducible ACE2 expression (Fig. S2A-B), and we infected these cells with closely related Omicron sub-lineages that differ in the presence of Spike mutation F486P, which increases ACE2 affinity ^30^. Using this approach, we found that increasing ACE2 expression augmented infection by Omicron sub-lineages, and this effect was more pronounced for lineages harboring the F486P polymorphism, XBB.1.5 and XBF, compared to those without it, such as XBB.1 (F486) and CH.1.1 (F486S) (Fig. S2C). While this result is consistent with the known role of ACE2 as the viral receptor for SARS-CoV-2, it contrasts with the observed in the V-ACE2^high^-TMPRSS2^high^ line, where overexpression of ACE2 attenuated TMPRSS2-dependent infection by Omicron sub-lineages. Together, these results suggest that high ACE2 levels may affect the ability of Omicron subvariants to utilize TMPRSS2 to mediate infection in some cellular contexts.

### Formation of ACE2/TMPRSS2 complexes reduces infection by Omicron but not pre-Omicron SARS-CoV-2 variants

It has been previously demonstrated that SARS-CoV-1 infectivity can be augmented by TMPRSS2-ACE2 complexes and that complex formation is promoted by the ACE2 CLD ^31^. This led us to investigate if SARS-CoV-2 entry is also regulated by the ACE2 CLD and its ability to recruit TMPRSS2. Since TMPRSS2 association with the ACE2 CLD prevents ACE2 shedding^30^, we screened the cell lines in Table I, for the presence of enzymatically active ACE2 in culture supernatants, as a marker for soluble ACE2. Detectable soluble ACE2 activity was observed across all engineered cell lines except the V-ACE2^high^TMPRSS2^high^ cell line, which showed high levels of ACE2 and TMPRSS2 at the cell surface (Fig. 2A-D), consistent with formation of ACE2-TMPRSS2 complexes that prevent ACE2 shedding.

**Figure 2.**
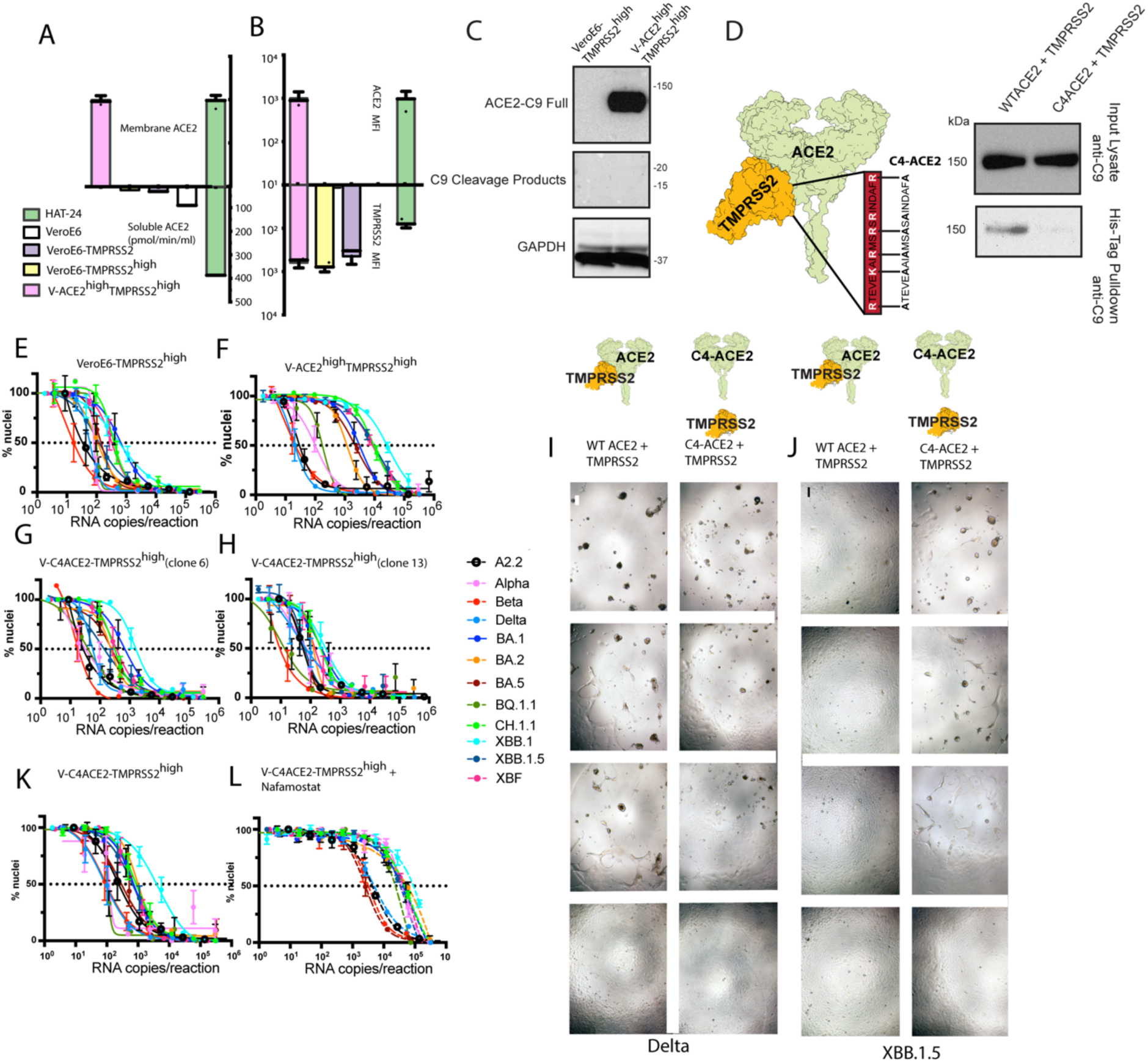
Abrogating TMPRSS2 interactions with ACE2 promotes replication in Omicron lineages **A.** Membrane and soluble ACE2 levels in engineered cell lines. Membrane ACE2 levels were measured by flow cytometry, mean fluorescence intensity is shown. The enzymatic activity of soluble ACE2 in the supernatant of 80% confluent cell lines after three days of culture was determined as previously described^32^. **B.** Ratios of cell surface ACE2 and TMPRSS2 detected via flow cytometry as per A. **C.** Expression of full length and cleaved ACE2 in VeroE6-TMPRSS2^high^ and V-ACE2^high^TMPRSS2^high^cells determined by immunoblotting with anti-C9 tag antibody. **D.** Left panel: ACE2 structure with known TMPRSS2 binding site in red (Structure PDB 6M17) and the sequence of the C4-ACE2 mutant to the right in black with alanine substitutions in bold. Right panel: TMPRSS2 pull-down of ACE2 WT and C4-ACE2 mutant, using an antibody against a His tag at the N-terminus of TMPRSS2 and blotting for ACE2 using an antibody against the C9 tag. **E-H.** Viral titration curves across 12 primary SARS-CoV-2 isolates with **E.** VeroE6-TMPRSS2^high^, **F.** V-ACE2^high^TMPRSS2^high^, **G.** V-C4-ACE2TMPRSS2^high^ **H.** V-C4ACE2TMPRSS2^high^. Input viral inoculate in E-H is presented here in RNA copies per infection/reaction**. I-J.** Contrasting replication across variants appearing in WT ACE2^high^, TMPRSS2^High^ and C4-ACE2^Mid^, TMPRSS2^High^ cell lines presented in F. and H., respectively, for pre-Omicron lineage I. Delta, and the Omicron lineage J. XBB.1.5. Fields of view from left to right are starting at 1/2000 dilution with 1/5 dilution steps in each successive frame. The Omicron lineage XBB.1.5 leads to extensive viral syncytia formation only in the presence of C4-ACE2, whilst the pre-Omicron Delta demonstrates viral syncytia at a similar level across WT and C4-ACE2 engineered cell lines. Scale bar is at 20 µM. K-L. Using the cell clone presented in H. we titrated virus in the **K.** absence or **L.** presence of saturating levels of the TMPRSS2 inhibitor Nafamostat. Data presented in K-L has been input as RNA copies of virus per variant.

In order to determine whether ACE2-TMPRSS2 complexes are present and contribute to the differential infection of cells co-expressing high levels of ACE2 and TMPRSS2, we turned to coprecipitation analyses and VeroE6-TMPRSS2**^high^** cells expressing either WT ACE2 or the C4-ACE2 mutant (Fig. S3). The mutations in C4-ACE2 remove a cluster of positive amino acid residues (Arg and Lys) in the ACE2 CLD known to be important for proteolytic regulation of ACE2 by both TMPRSS2 and ADAM17^31^. While coprecipitation experiments revealed the presence of complexes involving WT ACE2 and TMPRSS2, evidence of complex formation was not observed for CA-ACE2 and TMPRSS2 (Fig. 2D). Interestingly, all tested Omicron lineages infected C4-ACE2/TMPRSS2 co-expressing cells with much higher efficiency than cells co-expressing wild-type ACE2 and TMPRSS2 (Fig. 2E-H). These results suggest that the attenuation of Omicron sub-lineages imposed by the co-abundance of ACE2 and TMPRSS2 is dependent on the ACE2 CLD ability to recruit TMPRSS2. Indeed, while pre-Omicron variants like early Clade A and Delta support robust viral replication and syncytia formation in cells co-expressing WT ACE2 and TMPRSS2, comparable high titers and pronounced syncytia formation in Omicron subvariant cultures was only observed in cells that co-express C4-ACE2 but not WT ACE2 alongside TMPRSS2 (Fig. 2 I&J). Finally, we confirmed that infection mediated by C4-ACE2 remained TMPRSS2-dependent, employing the TMPRSS2 inhibitor Nafamostat. Indeed, substantial reductions in titers (shift from left to right) upon Nafamostat treatment demonstrated that replication of both pre-Omicron and Omicron subvariants was TMPRSS2-dependent (Fig. 2 L-M) when C4 ACE2 was present. In sum, these results support the concept that pre-Omicron variants can robustly use ACE2 and TMPRSS2 to mediate infection regardless of ACE2-TMPRSS2 complex formation, whilst Omicron subvariants replication is reduced in the presence of these complexes. Furthermore, uncoupling TMPRSS2 interactions with ACE2 by mutating the CLD removes this block and allows for robust TMPRSS2-dependent replication of Omicron subvariants.

### ACE2-TMPRSS2 complexes can facilitate viral entry of Omicron sub lineages but not viral spread

In order to confirm that pre-Omicron variants but not Omicron sub-lineages efficiently infect cells which allow for complex formation of ACE2 and TMPRSS2, we extended our analyses to every major variant detected during the 2020-2025 period. We found that infection by all pre-Omicron variants was consistently augmented (with the exception of Alpha) by the overexpression of WT ACE2 in the VeroE6-TMPRSS2^high^background. In contrast, infection of the resulting V-ACE2^high^TMPRSS2^high^ line by all tested Omicron sub-lineages was attenuated compared to the parental line. Interestingly, a gradient of increasing attenuation was observed, which broadly correlated with the time of subvariant emergence during the pandemic (Fig. 3A), with infection by contemporary Omicron sub-lineages being particularly inefficient (e.g. JN.1 sub-lineages). Finally, we confirmed that cells co-expressing C4-ACE2 and TMPRSS2 supported robust and comparable replication of all variants tested (Fig. 3A). In order to determine whether the attenuated infection of V-ACE2^high^-TMPRSS2^high^by the Omicron sub-lineages was the result of reduced viral entry, we employed SARS-CoV-2 spike pseudotyping.I nterestingly, pseudotyped particles bearing the S proteins of all previously tested variants were able to enter V-ACE2^high^-TMPRSS2^high^ cell lines with similar efficiency (Fig. 3B), regardless of whether WT ACE2 or C4-ACE2 was overexpressed. These findings suggested that the attenuated replication of Omicron sub-lineages in cells that co-express WT ACE2 and TMPRSS2 was not due to diminished viral entry but rather impairment of a post-entry step.

**Figure 3.**
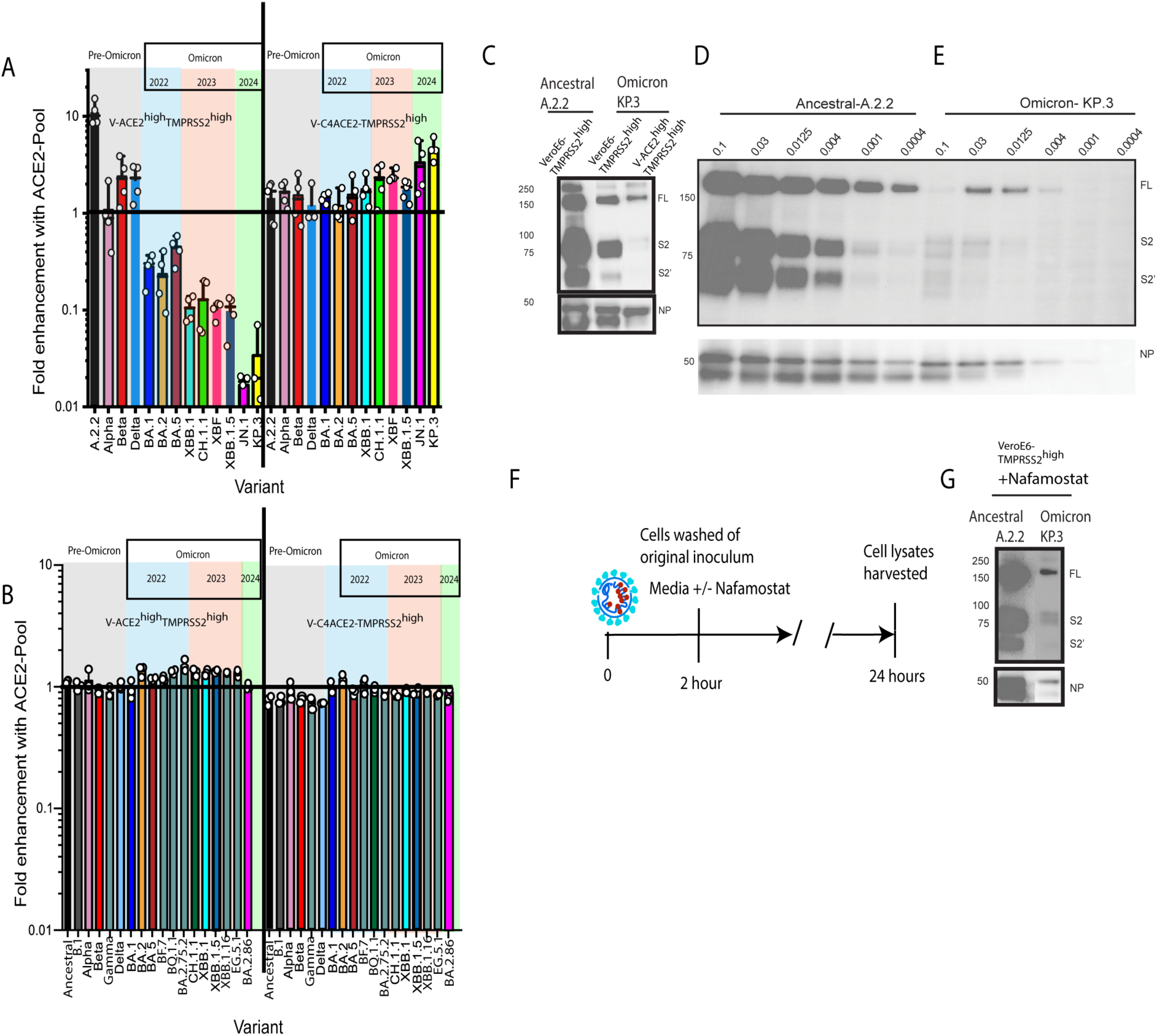
ACE2-TMPRSS2 complexes facilitate Omicron entry but not latter TMPRSS2-dependent spread **A&B.** Summary of fold augmentation and attenuation of all major variants to date across the two pools of ACE2 (WT ACE2 and C4-ACE2) when co-expressed with TMPRSS2 in a VeroE6 background. Fold attenuation or augmentation is calculated as the fold change of the engineered cell line expressing each ACE2 pool compared to the parental expressing TMPRSS2 and low ACE2 (VeroE6-TMPRSS2^high^). In **A.** each data points is presented from independent experiments. Data here represents replicating primary isolates over several rounds of infection. **B.** Pseudotyping of Spike can readily resolve single entry events. As in A. we used pseudotyping assays with cell co-expressing WT ACE2 and C4-ACE2 in addition to TMPRSS2. Fold increases/decreases are then compared to the parental VeroE6-T2 line which is set at the line 1. As in A. each data points is presented from independent experiments. **C.** Expression of Spike and Nucleocapsid proteins in the Omicron permissive VeroE6-TMPRSS2^high^ and non-permissive V-ACE2^high^TMPRSS2^high^ cell line following 24 hours of infection using the Omicron JN.1 sub-lineage KP.3. Ancestral clade A.2.2 is presented as a control. FL = Full length spike protein, S2 = S2 Spike only and S2’ = cleaved S2 spike. NP = Nucleocapsid protein. Position of protein molecular with standards in KDa are presented to the left of the panel. **D. & E.** As in C. Spike and Nucleocapsid proteins are presented following 24 hours of infection of the Omicron non-permissive cell line V-ACE2^high^TMPRSS2^high^. Here Spike and Nucleocapsid products of Ancestral (Clade A.2.2) and Omicron (KP.3) are presented across a range of viral multiplicities of infection that are indicated above each lane. **F.&G.** Here Omicron permissive cells (VeroE6-TMPRSS2^high^) are inoculated for 1 hour at an MOI of 0.025 with Ancestral and Omicron KP.3. Cells are washed thoroughly and then cultured in the presence of saturating levels of Nafamostat (10µM). Infection then proceeds for a 24-hour period, with Spike and Nucleocapsid proteins analysed as outlined in C.

Omicron sub-lineages have been reported to exhibit poor cleavage of the S protein at the S1/S2 cleavage site within infected cells, which may account for the reduced TMPRSS2 use during viral entry ^19,20^. To examine viral spike cleavage and spike-driven entry, we generated viral pseudotypes in the absence and presence of overexpressed ACE2 and ACE2-TMPRSS2. We observed robust cleavage of the S proteins of the Ancestral and Omicron sub-lineage across all conditions tested (Fig S5), suggesting that high expression of ACE2 and/or TMPRSS2 does not modulate S protein cleavage in pseudotype producing cells. We next examined infection of the VeroE6-TMPRSS2^high^ cell line (Omicron permissive) and the V-ACE2^high^TMPRSS2^high^ (Omicron attenuated) cell line with the KP.3 Omicron sub-lineage and the Ancestral Clade A.2.2 variant in order to determine whether cleavage of the viral S proteins in these cell lines is efficient. S protein fragments indicative of cleavage at the S1/S2 site were observed upon both Clade A.2.2 and KP.3 infection of the Omicron permissive VeroE6-TMPRSS2^high^ cells (Fig 3C). In contrast, spike expression and cleavage was reduced in KP.3 infected V-ACE2^high^TMPRSS2^high^ cells (Fig 3C). Further, no nucleocapsid cleavage was detected under these conditions (Fig 3C), consistent with lack of cytopathic effects that lead to nucleocapsid cleavage through activated caspases ^33^. These results are consistent with attenuation of Omicron subvariants in V-ACE2^high^TMPRSS2^high^ cells being due to inefficient S1/S2 cleavage. To resolve this further, we examined pre-Omicron clade A and Omicron KP.3 infection of non-permissive V-ACE2^high^TMPRSS2^high^ line across a range of inocula. Across both pre-Omicron and Omicron sub-lineages, lower levels of viral replication (i.e. decreasing viral inocula) were associated with lower cleavage of spike at the S1/S2 site (Fig 3D & E).

Given our findings that Omicron subvariant replication is attenuated in cells that form ACE2/TMPRSS2 complexes and that this outcome is likely dictated by a post-entry step, we next asked whether post-entry inhibition of TMPRSS2 would differentially affect pre-Omicron variants compared to Omicron sub-lineages. To this end, we added the TMPRSS2 inhibitor Nafamostat to the Omicron-permissive VeroE6-TMPRSS2^high^ cell line at 2 hours post-infection (Fig. 3F) and assessed viral replication by measuring cellular Spike protein levels at 24h post-infection. While we observed robust replication of the pre-Omicron Clade A lineage under these conditions, replication of the KP.3 Omicron sub-lineage was attenuated in a manner similar to that observed in the V-ACE2^high^TMPRSS2^high^ cell line (Fig. 3G). This suggests that while both pre-Omicron and Omicron variants are susceptible to complete TMPRSS2-inhibition, post-entry inhibition disproportionally attenuates Omicron lineages in a manner reminiscent to that observed in cells that readily form ACE2/TMPRSS2 complexes. This invites the interesting consideration that CLD-dependent association of ACE2 with TMPRSS2 may negatively affect Omicron sub-lineages’ ability to use TMPRSS2 at a post-entry step.

### ACE2 oligomerization with Solute Carriers SLC6A19 and SLC6A20 facilitates Omicron spread through increased TMPRSS2 surface activity and use

Apart from the well-known role of soluble ACE2 in the RAS pathway, ACE2 can also serve as a chaperone for the solute carrier SLC6A19 ^34,35^ and can act as an alternate chaperone for SLC6A20 in the absence of the primary chaperone Collectrin, which is an ACE2 homologue ^36–38^. Both solute carriers are sodium-dependent symporters of neutral amino acids that belong to the solute carrier 6 (SLC6) family and the subfamily of orphan transporters, comprised of SLC6A15-SLC6A20 ^39^. Both SLC6A19 and SLC6A29 have been reported to form a dimer of heterodimers with ACE2 and do so in an ACE2 CLD-dependent fashion, involving ACE2 amino acid residues 710 and 716 ^37,40^. Since this heterodimerization may sterically hinder ACE2 interactions with TMPRSS2 we hypothesized these solute carriers could influence Omicron sub-lineages infection in the tissues where they are expressed.

SLC6A19 is predominantly expressed in the small intestine and kidney, whereas SLC6A20 is mainly expressed in the small intestine, kidney, lungs, and brain ^41^. To address the pathophysiological relevance of these solute carriers for SARS-CoV-2 infection, we determined, at the single cell level, co-expression of ACE2 and TMPRSS2 with SLC6A19 and SLC6A20 in target tissues of SARS-CoV-2 infection, such as the small intestine (enterocytes), lung (type 2 pneumocytes) and the nasal cavity (ciliated nasal epithelia). We observed three cellular phenotypes of interest across these tissues. Firstly, ACE2^+^//SLC6A19^+^/TMPRSS2^+^ enterocytes. Secondly, low levels of ACE2^+^/Collectrin^+^/SLC6A20^+^/TMPRSS2^+^ type 2 pneumocytes. Thirdly, ACE2^+^/ SLC6A20^+^/TMPRSS2^+^ ciliated nasal epithelia (Fig. S4.). Given the potential for SLC6A19 and SLC6A20 to influence ACE2/TMPRSS2 dynamics and therefore viral replication, we generated clonal cell lines that expressed TMPRSS2 jointly with ACE2 and each solute carrier at equimolar levels using a bicistronic expression construct, resulting in the VeroE6 cell lines TASL-19 (**T**MPRSS2+**A**CE2+**SL**C6A**19**) and TASL-20 (**T**MPRSS2+**A**CE2+**SL**C6A**20**). Both cell lines expressed detectable surface levels of ACE2 and high surface expression of TMPRSS2 (Fig. S6), and the expression of SLC6A19 and SLC6A20 at the protein level was confirmed by western blotting of cell lysates (Fig. S6). In sum, we have generated cell lines that mimic ACE2, TMPRSS2 and solute carrier expression patterns found in viral target tissues.

We next analyzed the impact of SLC6A19 and SLC6A20 expression on ACE2 and TMPRSS2 interactions and permissiveness to SARS-CoV-2 infection. Although these solute carriers can bind to the CLD of ACE2, it is not known if this binding interferes with ACE2 interactions with TMPRSS2. To test this, we co-expressed TMPRSS2 with either ACE2 alone or ACE2 in conjunction with SLC6A19 and attempted to pull down ACE2 with TMPRSS2. While evidence of TMPRSS2/ACE2 complexes was readily observed in the absence of SLC6A19, co-expression of this solute carrier abolished complex formation (Fig. S6C.). Further, the Omicron sub-lineage KP.3 replicated in both TASL-19 and TASL-20 cell lines with higher efficiency as compared to the parental, highly permissive cell line VeroE6-TMPRSS2^high^ and induced extensive syncytia formation, which was not observed in the absence of solute carrier expression (Fig. 4A & B). Compared to the parental line, TASL-19 cells sustained 10-to 20-fold higher titers of every major SARS-CoV-2 variant (Fig. 4B), and we noted a trend towards higher titers for the more recent Omicron sub-lineages.

**Figure 4.**
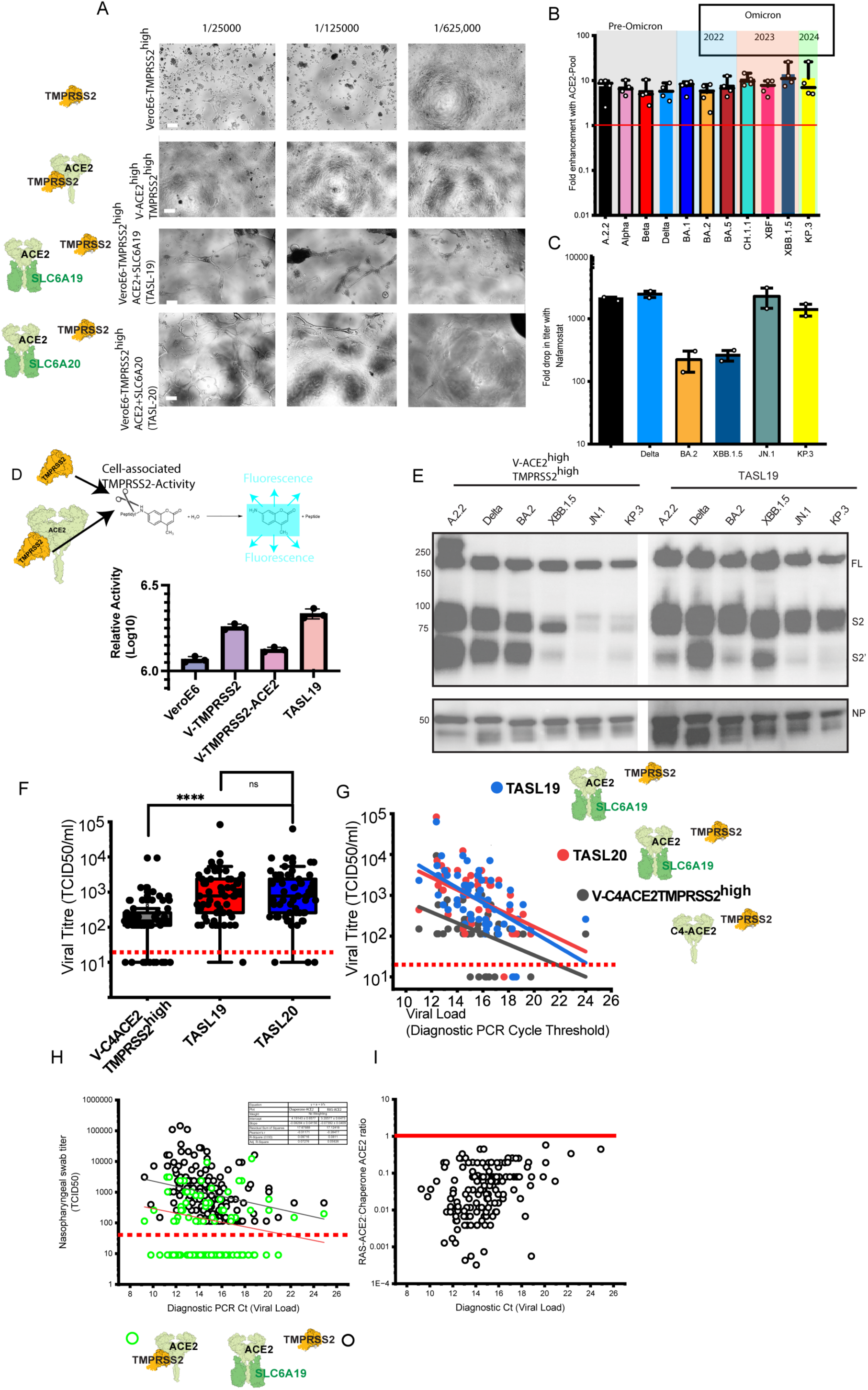
Solute carriers chaperoned by ACE2 augment infection across all SARS CoV-2 lineages. **A.** Cells were inoculated with the representative KP.3 Omicron isolate at dilutions stated above panels. Within 48 hours, extensive syncytia is observed primarily in TASL-19 and TASL-20 cell lines. Limited to no cytopathic effects are observed using the V-ACE2^high^-TMPRSS2^high^ cell line at equivalent viral inocula. Scale bars are at 50μm. **B.** Summary of fold augmentation of all major variants to date of the TASL-19 cell line. Fold augmentation is calculated as the fold change of the engineered TASL-19 line compared to the parental cell line expressing VeroE6-TMPRSS2^high^. In **B.** data points are presented from independent experiments. **C.** TMPRSS2 dependence in TASL-19 across representative pre-Omicron and Omicron lineages. Titers are calculated as outlined in Fig.1 D. with and without the TMPRSS2 inhibitor Nafamostat. Each data point represents independent experiments. **D.** Cell associated TMPRSS2 measured through cell culture of the Boc-Gln-Ala-Arg-AMC fluorogenic peptide for TMPRSS2 negative VeroE6, VeroE6-TMPRSS2^high^, V-ACE2^high^-TMPRSS2^high^ and the TASL-19 cell line. Each data point represents independent experiments. **E.** Analysis of viral Spike and Nucleocapsid proteins in Ancestral and Omicron KP.3 after 24 hours infection in **E. (**left panel), the Omicron non-permissive cell line V-ACE2^high^-TMPRSS2^high^ versus (right panel) the Omicron permissive TASL-19 cell line. For Spike, western blotting use the anti-S2 antibody reveals uncleaved Spike (FL), S2 and cleaved S2 (S2’). Molecular weight markers are indicated to the left of the figure and are in KDa. Western blot is representative of three independent expansions of pre-Omicron and Omicron lineages. **F. & G.** Utility of the TASL-19 and TASL-20 clones in primary viral isolation from clinical samples. Here 75 positive nasopharyngeal swabs with diagnostic PCR cycle thresholds less than 25 were collected and set aside at −80^0^C. Swabs were sterile filtered and then titrated on the V-C4ACE2TMPRSS2^high^, TASL-19 & TASL-29 cell lines. Of note the V-C4ACE2TMPRSS2^high^ line is already more sensitive in viral isolation compared to the parental VeroE6-TMPRSS2^high^ cell line and orders of magnitude more sensitive to the V-ACE2^high^-TMPRSS2^high^. **H. & I.** The consolidation of Chaperone ACE2 (TASL-19) versus angiotensin regulation sensitive RAS-ACE2 (V-ACE2^high^-TMPRSS2^high^) for use in SARS-CoV-2 phenotypic surveillance for primary nasopharyngeal swabs obtained in continued 2025 genotype to phenotype surveillance in Sydney NSW Australia. **H.** As. In H. viral titers from swabs are presented plotted against the diagnostic PCR cycle thresholds. In over two thirds of samples, no detectable titers were observed in RAS-ACE2 (V-ACE2^high^-TMPRSS2^high^) line and for presentation, titers are arbitrary designated a titer of 9. **I.** Here we define Omicron tropism as preferential infection of only cells expressing Chaperone-ACE2 (TASL-19). This is established by the ratio of titers of RAS-ACE2 (V-ACE2^high^-TMPRSS2^high^): Chaperone-ACE2 (TASL-19). Here Ancestral controls replicate equally with ratios approaching 1 designating a pre-Omicron tropism. As in H. RAS-ACE2 (V-ACE2^high^-TMPRSS2^high^) titers that are not detected are arbitrarily designated a titer of 9 to enable ratios to be calculated and presented.

To investigate the role of TMPRSS2 activity in TASL cells, we titered representative Omicron sub-lineages on the TASL-19 line in the presence of the TMPRSS2 inhibitor Nafamostat. This revealed large reductions in viral titers (Fig 4C), demonstrating that viral replication in TASL lines is TMPRSS2-dependent. Of interest across Omicron lineages this was more pronounced in contemporary JN.1 sub-lineages compared to earlier circulating Omicrons BA.2 and XBB.1.5. Further, to test if TMPRSS2 activity is differentially regulated by the co-expression of ACE2 alone or ACE2 co-expressed with SLC6A19, we incubated VeroE6 (TMPRSS2 negative), VeroE6-TMPRSS2^high^, V-ACE2^high^-TMPRSS2^high^ and the TASL-19 cell line in the presence of the TMPRSS2 Boc-Gln-Ala-Arg-AMC fluorogenic peptide substrate, which is not cell permeable (i.e. Resolves primarily surface TMPRSS2 activity in unpermeabilized live cells). Whilst *in situ* surface TMPRSS2 activity was detected on the VeroE6-TMPRSS2^high^ cell line, the V-ACE2^high^-TMPRSS2^high^ line revealed TMPRSS2 activity not significantly different to the TMPRSS2-negative VeroE6 control (Fig. 4D). This confirmed that overexpression of ACE2 not only reduces Omicron sub-lineages ability to utilize TMPRSS2, but it also inhibits surface TMPRSS2 enzymatic activity. In contrast, co-expression of ACE2 with SLC6A19 in TASL-19 cells led to similar TMPRSS2 surface activity as in VeroE6-TMPRSS2^high^ cells (Fig. 4D), indicating that the presence of this solute carrier can rescue ACE2-dependent inhibition of TMPRSS2. Therefore, we conclude that expression of the solute carriers SLC6A19 or SLCA20 can dissociate inhibitory ACE2-TMPRSS2 complexes, increasing TMPRSS2 activity at the cell surface. Overall, our findings suggest that Omicron subvariants depend on TMPRSS2 activity at the cell surface, which can be negatively regulated by formation of ACE2-TMPRSS2 interactions. In contrast pre-Omicron subvariants can utilize TMPRSS2 to drive infection and syncytia formation, regardless of ACE2-TMPRSS2 complex formation.

To further analyse the differential viral replication observed in V-ACE2^high^-TMPRSS2^high^ cells (non-permissive to Omicron lineages) and TASL cells (highly permissive to Omicron infection), we measured the accumulation of the viral proteins Spike and Nucleocapid following 24 hours post infection. Whilst the successive appearance of key Omicron lineages is associated with decreasing spike expression and lack of S1/S2 spike cleavage in the V-ACE2^high^-TMPRSS2^high^ line (Fig. 4E), comparable levels of spike expression and S1/S2 cleavage were observed with all Omicron lineages in the TASL-19 line (Fig. 4F.). These findings reinforce that solute carriers such as SLC6A19 and SLC6A20 can rescue TMPRSS2 activity and restore permissiveness to Omicron sub-lineage infection in cells that co-express ACE2 and TMPRSS2.

### Cells co-expressing SL6A19 alongside ACE2 and TMPRSS2 are highly permissive to infection and can be used for sensitive isolation of primary isolates

Isolation of primary isolates for phenotypic testing still represents the most stringent means to enable testing of neutralization responses following vaccination and also for continued surveillance and testing in therapeutic trials. As part of the NIH sponsored OTAC clinical trial, testing of emerging variants using novel hyper-immune globulin formations, made isolation of contemporary variants in a manner that led to rapid expansion of high titer viral preparations paramount in sustaining screening activities and testing of forumulations for this trial. To test the utility of the both TASL cell lines in this and other diagnostic settings, we compared these cells with the highly permissive the V-C4ACE2TMPRSS2^high^ line (Fig. 4F. & G.) for virus isolation. For this, we analyzed SARS-CoV-2 isolation from 75 diagnostic samples derived from a SARS-CoV-2 genotype to phenotype surveillance program ^26^. We measured isolation rates of 58% using the V-C4ACE2TMPRSS2^high^ cell line versus 76% and 77% for the TASL-19 and TASL-20 cell line, respectively (Fig. 4F. & G.). To establish a process to resolve pre-Omicron versus Omicron tropism in real-time, we we titered over 200 nasopharyngeal swabs across the V-ACE2^high^-TMPRSS2^high^ line (representative of RAS-ACE2) and TASL-19 (representative of chaperone ACE2). In an era of Omicron JN.1 sub-lineages, significant reductions in titers were predicted to be observed in cells expressing RAS-ACE2 relative to chaperone ACE2). Whilst 156 swabs were positive after 3 days culture in the TASL-19 line, only 48 were positive in the V-ACE2^high^-TMPRSS2^high^ line and all viruses attained lower titers than in the TASL-19 line as reflected in the RAS-ACE2:Chaperone ACE2 ratios (Fig. 4I).

## Discussion

Resolution of viral entry pathways is essential to track the fitness of viral variants and their ability to spread, and to understand the pathogenesis and potential clinical manifestations as they appear in the community. In most instances the shifts in viral entry are subtle, however there are occasions where a significant accumulation of changes in a viral glycoprotein can enable a step-wise shifts in how the virus engages and infects cells and tissues. Whilst most SARS-CoV-2 infections are rapidly cleared, prolonged viral replication in immunocompromised or immunosuppressed patients has been shown to lead to an accumulation of amino acid substitutions or deletions within Spike ^42–45^. This accumulation of mutations is the primary hypothesis for the emergence of new immune-escape variants, such as the VOC Omicron, which is the parent of all currently circulating variants ^44^. Omicron has evolved to target cells and tissues in a manner different from earlier circulating lineages. From preliminary studies by ourselves and others, it was immediately apparent that Omicron lineages were engaging the serine protease TMPRSS2 differently than Delta and earlier pre-Omicron variants ^17,19,20^. This change in entry manifested *in vivo* in animal models ^14^, as well as in a marked decrease in COVID-19-associated pneumonia prevalence in patients infected with Omicron ^46^. Ironically, the more transmissible the virus became in humans, the more attenuated it was observed to be *in vitro* where entry factors ACE2 and TMPRSS2 were co-expressed. Herein, using extensive panels of primary viral isolates from early 2020 through to 2025, we have mechanistically dissected the evolving entry requirements of SARS-CoV-2 variants across the pandemic to date. The key to resolving this dichotomy between *in vitro* and *in vivo* observations was to build beyond the known SARS-CoV-2 receptor complex of ACE2 and TMPRSS2 and bring in two tissue specific solute carriers, SLC6A19 and SLC6A20, both of which are known to form dimers of heterodimers with ACE2^37,47^. In the latter setting we reveal a new paradigm for SARS CoV-2 entry. Here pre-Omicron SARS-CoV-2 can readily utilize distinct ACE2 pools, whether in association with TMPRSS2 (hereon referred to as RAS-ACE2) or in association with a solute carrier (hereon referred to as chaperone ACE2). In contrast, we found Omicron variants to only replicate efficiently in chaperone ACE2 pool environments. Mechanistically, this translates to a requirement for sufficient TMPRSS2 activity at the cell surface. Given this activity is negatively regulated by coexpression of ACE2, attenuation in Omicron replication is observed in the RAS-ACE2 pool setting. However, if ACE2-interacting solute carriers are expressed, the resulting complexes with ACE2 prevent ACE2-TMPRSS2 interactions and as such rescue both TMPRSS2 surface activity and viral replication.

The initial phenotypic shift in entry for SARS-CoV-2 primarily took place from Delta to Omicron BA.1 and BA.2. The latter has been studied extensively and the changing requirements for entry have been documented and proposed using a variety of *in vitro* cell lines and models. The Omicron entry hypothesis is broadly based on the inefficient cleavage of the S1/S2 Spike domains by Furin, resulting in increased non-cleaved S1/S2, and reduced fusogenic activation of S2 by TMPRSS2 *trans* cleavage and as a direct consequence increased endosomal entry. In the latter endosomal entry pathway, Spike utilizes the cysteine protease Cathepsin L to activate and drive S2 fusion across the endosomal membrane ^19,20^ which does not require prior S1/S2 Spike cleavage. Whilst many *in vitro* studies support this hypothesis, studies with primary cell cultures and replication of Omicron lineages *in vivo* ^5,48^ reveal a continued requirement for TMPRSS2-mediated *trans* cleavage for S2 fusogenic activation. Our study herein mechanistically resolves the complexities of observations initially observed *in vitro* with engineered cell lines to that observed in primary cell models. Here we outline this at several levels. Firstly, Omicron lineages may indeed be forced into using the endosomal pathway for viral replication when surface TMPRSS2 activity is blocked in the presence of RAS-ACE2. Whilst the lack of detectable S1/S2 Spike cleavage products can be associated with reduced furin or TMPRSS2 processing, it is also observed when viral replication is limiting, e.g. in Omicron cultures attenuated by RAS-ACE2 or in pre-Omicron cultures seeded with low multiplicity of infection. Indeed, detection of S1/S2 cleavage products can be enhanced by increasing the initial inoculum in both pre-Omicron and Omicron cultures. Therefore, we would conclude that the Furin Cleavage Site in Omicron lineages is sustained and that lack of S1/S2 detectable spike cleavage in Omicron cultures is a feature of low levels of viral replication as a consequence of another activity that drives attenuation. This is consistent with observations from direct biochemical assays that quantitatively measured furin cleavage efficiency and found Omicron Spike sequences to have an increased intrinsic cleavability by furin compared to earlier variants ^49^, despite some cell-based data suggesting otherwise. Furthermore, when attenuated Omicron replication is rescued by the presence of chaperone-ACE2, robust S1/S2 cleavage is also observed and thus further consistent with S1/S2 cleavage appearing during efficient viral replication. In addition to driving greater levels of virus replication, chaperone ACE2 and TMPRSS2 also enabled extensive cell to cell fusion in Omicron cultures and thus points to cell-cell spread being promoted in this setting. For pre-Omicron models syncytia formation was observed to be dependent on the spatial orientation of TMPRSS2 in relation to the S protein, i.e. TMPRSS2 needed to be present in the opposing cell membrane to activate S protein and induce cell-cell fusion ^50^. Our observations therefore support a model where the spatial orientation of TMPRSS2 complexed to ACE2 prevents Omicron Spike engagement in cell-cell spread & associated cell-cell fusion. However, when TMPRSS2 is excluded from complexing with ACE2 by the latter’s association with solute carriers SLC6A19 and SLC6A20, cell-cell spread/fusion of Omicron is favoured, resulting in high levels of replication and syncytia formation. In contrast, pre-Omicron variants are promiscuous in their use of TMPRSS2 and can support cell-cell spread both in the presence and absence of TMPRSS2-ACE2 complexes.

Whilst the above ACE2 pool model explains the phenotypic shift of the virus over the pandemic, there are two hypotheses that support the manifestation of this trajectory over time. Firstly, the virus has simply evolved to target the upper respiratory tract better, with the step-wise accumulation of Spike mutations that increase tropism for these tissues, given the resulting transmissibility advantage. Alternatively, the appearance of lineages with significant changes in Spike may be a feature of unknown chronic infections in other tissue reservoirs that harbor permissive ACE2 pools similar to those found in the upper respiratory tract. Here enterocytes across the intestine would represent the most obvious target as this is where ACE2 is the most abundant *in vivo* and would provide a rich environment of cellular substrates for the virus to replicate in. This setting is similar to that outlined for HIV-1, where cellular substrate availability in mucosal tissue dictates *in vivo* infection outcomes^51^. By replicating in abundant cellar targets, SARS-CoV-2 may have simply consolidated to this abundant substrate within the intestinal tract. This is consistent with continued observations of SARS-CoV-2 in waste-water viral loads (https://data.who.int/dashboards/covid19/wastewater). Furthermore, both SCL6A19 and SLCA20 are most abundantly expressed in the gut, which further supports a role for chaperone ACE2 pools in supporting viral replication. However, whilst SLC6A19 is primarily restricted to enterocytes, SLC6A20 is also expressed throughout the respiratory tract and on known SARS-CoV-2 target cells. Whilst ACE2 is the primary chaperone for SLCA19, the ACE2 homologue Collectrin can also act as a chaperone for SLC6A20. Therefore, the relative abundance of ACE2 and Collectrin will likely dictate which chaperone forms a complex with SLC6A20 in different tissues. Whilst ACE2 has a gradient of decreasing expression from the upper to lower respiratory tract ^52^, collectrin expression in the respiratory tract is limited. Interestingly, SLC6A20 has been identified in several genome-wide association studies (GWAS) as having one of the strongest associations with COVID-19 susceptibility ^53^. Furthermore, tight complexes of SLC6A19 and SLC6A20 with ACE2-RBD have been resolved in cryo-EM studies ^37^, and both solute carriers have been independently suggested as potential targets in the treatment of COVID-19 ^54,55^. Overall, the evidence presented by ourselves and others suggests an important role of these solute carriers in COVID-19 pathophysiology, regulation of tissue specific viral tropism and importantly susceptibility to infection.

While the preferential utilization of chaperone ACE2 pools may represent one of the key changes reflecting the evolving tropism of of SARS-CoV-2 Omicron infection, it is likely not the only factor. Structural stability of the Spike trimer ^56^, the closed/open prevalence of the receptor binding domain ^57–61^, ACE2 affinity ^62,63^, ability to navigating pre-existing neutralizing antibodies ^17,19,20,26,64,65^, binding to heparan sulfates ^66^, efficient S1/S2 spike cleavage ^10,67^ and resistance to human airway trypsin (HAT) ^68^ may all contribute to a highly transmissible SARS-CoV-2 Omicron phenotype. Of note, all dominant lineages to date have a competitive advantage that is inversely related to use of the RAS-ACE2 pool. i.e. The greater levels of attenuation in the RAS-ACE2-TMPRSS2 pool, the greater advantage in viral transmission. In pre-Omicron variants that became variants of concern (e.g. Alpha and Delta) changes at position 681 in the furin cleavage site resulted in significant growth advantage in the community ^69,70^ but replicated at lower levels *in vitro* compared to Ancestral virus in RAS-ACE2-TMPRSS2 environments. The same is true with respect to the unknown competitive advantage of BA.2.86 and JN.1 sub-lineages over XBB.1 derived lineages. Here, the change at 681 from H to R again led to a competitive advantage in the community in addition to a significant shift in RAS-ACE2-TMPRSS2 attenuation. The change at the 681 site is associated with greater furin cleavage ^10^ and increased resistance to cleavage by human airway trypsin-like protease (HAT). The culmination of this change may benefit from both known fitness gains but at the opportunity cost of tropism promiscuity, due to reduced ability to sustain replication in RAS-ACE2-TMPRSS2 pools. Curiously, Ancestral clade A remains the most competitive lineage in RAS-ACE2-TMPRSS2 environment and this may represent a competitive transmission gain in another animal reservoir. Of note, once in the human population, there has been a clear trajectory for the virus to increasingly favour chaperone ACE2 pools and this is consistent with the consolidation of Omicron sub-lineage infection to the upper respiratory tract.

As the high level of spread of SARS-CoV-2 globally continues and is compounded by the risk of co-circulation in other hosts, phenotypic surveillance of variants is needed to determine the consequences of continuing viral evolution. The impact and result of chronic infection in humans and/or in other animal reservoirs is still unknown and needs to be monitored carefully. Rapid detection of shifts in viral tissue tropism and linkage to severe clinical presentation are important in real-tracking many viruses including SARS-CoV-2. However, to enable detection of tropism shifts, mechanistic resolution of the changing entry requirements is needed to distill this knowledge into entry screening platforms. Herein we have determined a key change in Omicron subvariant’s infection leading to differential replication in cellular environments with distinct ACE2 pools. We also outlined assays based on clonal cell lines that can readily determine how an emerging variant benefits from distinct ACE2 pools and importantly if it is on a trajectory towards or away from pre-Omicron tropism features associated with more severe disease. Rapid phenotypic surveillance of future variants of interest or other ACE2 using coronaviruses, may be key in identifying those which have higher clinical severity scores or tissue-specific clinical manifestations such as lung-related pneumonia. Recent studies have observed the Bat-derived merbecovirus HKU5-CoV lineage 2 to replicate in human respiratory and intestinal cultures ^64^ highlighting the need for continued phenotypic surveillance beyond SARS-CoV-2 to gauge the pandemic potential following zoonosis.

This work further highlights the need to understand ACE2 pools at greater resolution *in viv*o and determine how the role of ACE2 as a chaperone for solute carrier proteins may be driving the evolution of SARS-CoV-2 cellular tropism. At a minimum, this work enables platforms *in vitro* that can increase sensitivity of SARS-CoV-2 isolation, detection and phenotypic monitoring of all SARS CoV-2 variants. This will have a significant impact in diagnostics and future surveillance and management of other ACE2 using coronaviruses with pandemic potential. Importantly, these tools can now be used in rapid isolation and phenotypic characterization of SARS-CoV-2 lineages spreading within the community where viral genotypes and clinical severity are continuing to be studied and linked.

## Methods

### Cell culture

HEK293T cells (thermo Fisher, R70007), HEK293T derivatives including HAT-24 ^17^, VeroE6-TMPRSS2 (CellBank Australia, JCRB1819), VeroE6-TMPRSS2^High^ ^28^, V-ACE2^high^-TMPRSS2^high^, V-C4ACE2TMPRSS2^high^, V-TMPRSS2-ACE2+SLC6A19 (TASL-19) and V-TMPRSS2-ACE2+SLC6A20 (TASL-20) cell lines were cultured in Dulbecco’s Modified Eagle Medium (DMEM; Gibco, 11995073) with 10% fetal bovine serum (FBS) (Sigma; F7524). HEK293T-TetOne-ACE2 cells were cultured in DMEM with 10% fetal bovine serum (FBS) with 0.05µg/mL puromycin (Sigma; P8833). VeroE6 cells (ATCC CRL-1586) were cultured in Minimum Essential Medium (Gibco; 10370-021) with 10% FBS, 5% penicillin-streptomycin (Gibco; 15140122) and 1% L-glutamine (Gibco; 25030-081). Calu3 cells were maintained in DMEM/F12 media (Gibco;12634010) supplemented with 1% MEM non-essential amino acids (Sigma, M7145), 1% Glutamax (Gibco; 35050061) and 10% FBS. All cells used herein were only cultured and used within a range of 20 passages. Each cell line was tested and found negative for mycoplasma (Mycoplasma Testing Facility, UNSW).

### Generation of stable cell lines

For generating VeroE6-TMPRSS2^High^ cells that stably express mutant C4-ACE2, expression plasmid pRRLsinPPT.CMV.GFP.WPRE ^71^ was first modified to carry a multiple cloning site (MCS) at the 3’ end of GFP open reading frame (ORF) using Age1 and Sal1 cut sites. The mutant form of hACE2 (C4-ACE2) carrying mutations described previously by Heurich et al ^31^was synthesized as a synthetic gBlock (IDT) and shuttled into the above plasmid using Xba1/Xho1 cut sites thus replacing the GFP ORF with C4-ACE2.

For generating HEK293T-TetOne-ACE2 cells, expression plasmid pLVX Tet-One Puro (Clontech) was modified to have a MCS at the 3’end of Tet responsive promoter TRE3GS using EcoR1/BamH1 cut sites. ACE2 ORF was amplified from pRRLsinPPT.CMV.ACE2.WPRE plasmid ^17^ and shuttled into pLVX Tet-One Puro-MCS expression plasmid using Not1/Xho1 sites to generate into pLVX Tet-One Puro-ACE2 plasmid. Plasmid sequences were validated by Nanopore Sequencing with the Rapid Barcoding Kit 96 (Oxford Nanopore Technologies) using the manufacturer’s protocol. The sequencing data was exported as FASTQ files and analyzed using Geneious Prime (v22.2). Open reading frames were then verified using Sanger Sequencing.

For generating V-TMPRSS2-ACE2+SLC6A19 (TASL-19) and V-TMPRSS2-ACE2+SLC6A20 (TASL-20) cell lines, expression plasmid pRRLsinPPT.CMV.ACE2.WPRE ^71^ was first modified to remove the stop codon at 3’end of ACE2 orf. A synthetic gblock with P2A and a MCS carrying Nhe1 cut site was then synthesized (IDT) and inserted into the above plasmid at the 3’end of ACE2 to generate ppt-ACE2 MCS plasmid. SLC6A19 (Addgene #161397) and SLC6A20 (Addgene#161409) orfs were amplified with forward and reverse primers containing Nhe1 and Xho1 cut sites respectively. The amplicons were inserted into ppt-ACE2 MCS plasmid using Nhe1/Xho1 cut sites to generate bicistronic plasmids pptACE2-P2A-SLC6A19 and pptACE2-P2A-SLC6A20. Plasmid sequences were validated by Nanopore Sequencing with the Rapid Barcoding Kit 96 (Oxford Nanopore Technologies) using the manufacturer’s protocol.

Cells expressing C4-ACE2, inducible ACE2 and bicistronic ACE2 and solute carriers were generated by lentiviral transductions as previously described ^17^. Briefly, lentiviral particles expressing the above genes were produced by co-transfecting expression plasmids individually with a 2nd generation lentiviral packaging construct psPAX2 (courtesy of Dr Didier Trono through NIH AIDS repository) and VSVG plasmid pMD2.G (Addgene, 2259) in HEK293T producer cells using polyethyleneimine as previously described ^72^.Virus supernatant was collected 72 h post-transfection, pre-cleared of cellular debris and centrifuged at 28,000 × g for 90 min at 4°C to generate concentrated virus stocks. Lentiviral transductions were then performed on VeroE6-TMPRSS2^High^ cells to generate V-C4ACE2TMPRSS2^high^, TASL-19, TASL-20 and HEK293T cells to generate HEK293T-TetOne-ACE2. The expression of ACE2 in the latter was induced with 200 ng/ml of Doxycycline (Sigma, D9891) as per the manufacturer’s instructions (ClonTech). Screening of clones was based on expression of high levels of ACE2 following Doxycycline in addition to being highly susceptible to infection with the early clade SARS-CoV-2 isolate A.2.2. HEK293T-TetOne-ACE2 cells were further modified to express hTMPRSS2 ^28^ by lentiviral transductions to generate HEK293T-TetOne-ACE2-TMPRSS2 cells.

### ACE2 Affinofile Assay

Cells, engineered with Doxycycline induced ACE2, were trypsinised and seeded as suspension at 5 × 10^3^ cells/well in 384-well plates. Cells were exposed to varying amounts of Doxycycline for a minimum of four hours and at then inoculated with SARS-CoV2 variants for 72 hours. Following this, cells were stained live with 5% v/v nuclear dye (Invitrogen, R37605) and whole-well nuclei counts were determined with an INCell Analyzer 2500HS high-content microscope and IN Carta analysis software (Cytiva, USA). Data was normalized to generate sigmoidal dose–response curves (average counts for mock-infected controls = 100%, and average counts for highest viral concentration = 0%) and median Virus Effective (VE50) value for each doxycycline concentration was obtained with GraphPad Prism software ^27^

### Generation of pseudovirus particles and cell entry studies

The generation of vesicular stomatitis virus-based pseudovirus particles bearing SARS-CoV-2 S proteins and cell entry studies were conducted in accordance to previously published protocol ^73^. At 24 h post transfection, 293T cells expressing the respective S protein, VSV-G or dsRed (negative control) were inoculated with VSV-G-transcomplemented VSV∗ΔG(FLuc) (kindly provided by Gert Zimmer)^25^ and incubated for 1 h at 37°C and 5% CO2. Subsequently, the inoculum was aspirated and the cells were washed with phosphate-buffered saline (PBS), before cell culture medium with anti-VSV-G antibody (culture supernatant from I1-hybridoma cells; ATCC no. CRL-2700) was added. Of note, cells expressing VSV-G received cell culture medium without antibody. At 16–18 h post inoculation, the culture supernatants were collected, centrifuged to remove cellular debris (4,000 × g, 10 min), and clarified supernatants were aliquoted and stored at −80°C until further use. For cell entry studies, target cells were grown in 96-well plates to 50–90% confluence before they were inoculated with identical volumes of the respective pseudovirus particles and incubated for 16–18 h at 37°C and 5% CO2. Then, cell entry efficiency was assessed through measurement of virus-encoded luciferase activity in cell lysates. Cells were incubated for 30 min at room temperature with PBS containing 0.5% Tergitol (Carl Roth). Next, lysates were transferred to white 96-well plates, mixed with luciferase substrate (Beetle-Juice, PJK), and luminescence was measured with a Hidex Sense plate luminometer (Hidex).

### Viral isolation, propagation and titration from primary specimens

SARS-CoV-2 variants were isolated from diagnostic respiratory specimens as previously described and through the primary diagnostic provider Douglas Hanly Moir under UNSW ethics IREC5100 ^27^. Briefly, specimens testing positive for SARS-CoV-2 (RT-qPCR, Seegene Allplex SARS-CoV-2) were sterile-filtered through 0.22 µm column-filters at 10,000 x *g* and serially diluted (1:3) on C4-VeroE6-T2 or TASL-20 cells (5 x 10**^3^** cells/well in 384-well plates) and incubated for 72 hours. Upon confirmation of cytopathic effect by light microscopy, a minimum of 80ul of culture supernatant were added initially to a pellet of TASL-20 cells (1.0 × 10**^6^** cells) for 30 minutes and then subsequently transfer to a 6-well plate (well with 2 mL of MEM-2%FBS final) and incubated for 24 to 48 h (or until cytopathic effects had led to loss of >50% of the cell monolayer). The supernatant was cleared by centrifugation (2000 x *g* for 10 minutes), frozen at −80°C (passage 2), then thawed and titrated to determine median 50% Tissue Culture Infectious Dose (TCID50/mL) on VeroE6-TMPRSS2^High^ cells according to the Spearman-Karber method ^74^. Viral stocks used in this study correspond to passage 3 virus, which were generated by infecting VeroE6-TMPRSS2^High^ cells at MOI=0.025 and incubating for 24 h before collecting, clearing, and freezing the supernatant as above in 100 µl aliquots. Sequence identity and integrity were confirmed for both passage 1 and passage 3 virus via whole-genome viral sequencing using an amplicon-based Illumina sequencing approach, as previously described ^75^. The latter was also used in parallel for sequencing of primary nasopharyngeal swabs.

Virus titrations were carried out by serially diluting virus stocks (1:5) in MEM-2%FBS, mixing with cells initially in suspension at 5 × 10^3^ cells/well in 384-well plates and then further incubating for 72 h. The cells were then stained live with 5% v/v nuclear dye (Invitrogen, R37605) and whole-well nuclei counts were determined with an INCell Analyzer 2500HS high-content microscope and IN Carta analysis software (Cytiva, USA). Data was normalized to generate sigmoidal dose–response curves (average counts for mock-infected controls = 100%, and average counts for highest viral concentration = 0%) and median Virus Effective (VE50) values were obtained with GraphPad Prism software ^27^. To assess the TMPRSS2 usage of the virus isolates, titrations on VeroE6-TMPRSS2^High^ were performed in the presence of saturating amounts of Nafamostat (20 μM). Titrations performed in parallel in equivalent volumes of DMSO served as controls and were used to calculate fold drops in VE50.

### Phenotyping of ACE2 and TMPRSS2 on cell lines

Cell lines (2 x 10^5^ cells) were labelled with phycoerythrin-conjugated TMPRSS2 (Clone: S20014A, BioLegend) and AlexaFluor 647-conjugated ACE2 (Clone: 535919, R&D Systems) for 30 mins on ice in the dark. Cells were washed once with FACS wash (phosphate buffered saline containing 2 mM EDTA and 1% heat-inactivated foetal bovine serum), prior to fixation with 1% paraformaldehyde (final) and acquisition by a BD LSRFortessa^TM^ flow cytometer (BD Biosciences). Flow cytometry analysis was performed using FlowJo analysis software (version 10.8.0, BD Biosciences).

### Detection of catalytic ACE2 in cell supernatants

Soluble ACE2 activity in culture supernatants was measured as previously described ^32^.

#### Single resolution of known SARS-CoV-2 entry factors with ACE2 associated solute protein carriers

Single cell RNAseq data from nasal ^76^ and ileum ^77^ were obtained from covid19cellatlas.org, while lung data ^78^ was obtained from GEO (GSE171524). These data were analyzed with Seurat V4.3.0 as described previously ^79^. Their accompanying metadata, which includes information such as sample ID, sample status, and cluster annotations (cell types), were added to Seurat objects using the “AddMetaData” function. Read counts were normalized using SCTransform, before reanalysis with the standard Seurat workflow of “RunPCA,” “FindNeighbours,” “FindClusters,” and “RunUMAP.” Cluster identities were assigned using published cluster annotations and plots were generated with “DimPlot”, “Featureplot” and “DotPlot”, to illustrate expression of *ACE2*, *TMPRSS2*, *SLC6A14*, *SLC6A19* and *SLC6A20*.

#### Expansion and Purification of SARS-CoV-2 Viral Products

Cells (VeroE6, VeroE6-TMPRSS2^High^, V-ACE2^high^-TMPRSS2^high^, TASL19 and TASL20) were pelleted at 1×10^6^ cells and then resuspend each respective cell type pellet at MOI=0.025 (based on the TCID50 of previous stock titration for VeroE6-TMPRSS2^High^). Viral media/cells mixture were incubated at 37 C for 1 h. Exposed cells were washed twice MEM media supplemented with 2% Fetal Calf Serum (v/v) (MEM-2% media). After the final wash, infected cell pellets were resuspended in 2 mL MEM-2% media and were added to a 6-well plate. Plated infected cells were incubated at 37 C for 1 h. For Nafamostat treatment, media was removed and replaced with media containing Nafamostat at 10μm final concentration. All cells were then incubated at 37 C for 24 hr prior to subsequent harvesting of supernatant or cells.

Infected cells were harvested by washing once in 2 mL PBS, then dissociation from 6 well-plates using 1 mL TrypLE per well. Cells were incubated at 37 C for 3 min, then was neutralised with 2 mL MEM-2% media and collected into 2 ml conical tubes. Cells were centrifuged at 3000xrpm for 3 min, then supernatant was removed, and cells were resuspended in 1 mL of PBS. PBS wash step was repeated twice. After final wash, supernatant was removed, and cell pellets were kept and stored at −80 C until use.

#### Western Blotting of SARS-CoV-2 Spike Cleavage

TCA concentrated viral proteins were resuspended in NuPAGE LDS Sample Buffer (4x) and NuPAGE Sample Reducing Agent (10X) (Thermo Fisher), then samples were boiled at 100°C for 10 mins. Infected cell pellets were lysed in RIPA buffer (Thermo Fisher) with EDTA-free Protease Inhibitors (Roche), which was then diluted in NuPAGE LDS Sample Buffer (4x) and NuPAGE Sample Reducing Agent (10X), then samples were boiled at 100°C for 10 mins. All samples underwent electrophoresis using a NuPAGE Bis-Tris Protein Gel (Thermo Fisher), then gel separated proteins underwent a wet transfer. Transfer membranes were incubated with SARS/SARS-CoV-2 Spike Protein S2 antibody (1:2000, MA535946, ThermoFisher), anti-GAPDH antibody (1:4000, abcam, ab8245) or anti-SARS-CoV-2 Nucleoprotein (1:2000, Cellabs), then were probed with donkey anti-mouse HRP conjugate (1:5000, A16011, ThermoFisher). Visualization was performed using a G:Box (Syngene).

#### His-Tag Protein Pulldown

Gene-constructs of WT=ACE2, C4-ACE2 mutant, TMPRSS2 and Bicistronic ACE2:SLC6A19, that were described previously, were subcloned cloned into the pCDNA6 expression vector using cut site enzymes XbaI and XhoI. ACE2 constructs were modified to express a C9-tag on the C-terminal. The TMPRSS2 gene construct was modified to express a 6xhis-tag on its C-terminal end. As above all pCDNA6 plasmid constructs of all genes were verified using Nanopore Sequencing with the Rapid Barcoding Kit 96 (Oxford Nanopore Technologies) using the manufacturer’s protocol. pCDNA6-WTACE, pCDNA6-C4-ACE2 and pCDNA6-Bicistronic ACE2:SLC6A19 were each transiently transfected using PEI into HEK293-T cells with pCDNA6-TMPRSS2-histag as previously described ^71^. Transient transfections used a total of 5μg DNA with 5 million cells to ensure there was enough material for later pulldown experiments. The ACE2:TMPRSS2 DNA ratio was 2:1, with empty pCDNA6 being used as a DNA filler. Transfected cells incubated at 37 C for 48 hrs. After incubation, cells were harvested and washed with PBS.

Transfected cells were lysed in RIPA buffer (Thermo Fisher) with EDTA-free Protease Inhibitors (Roche). Lysate was applied on Ni-NTA Magnetic Beads (S1423S, New England Biolabs) using manufacturers protocol. A fraction of raw lysate was saved for blotting. After samples were eluted from the magnetic beads, they were diluted in NuPAGE LDS Sample Buffer (4x) and NuPAGE Sample Reducing Agent (10X) (Thermo Fisher), then samples were boiled at 100°C for 5 mins. Samples underwent SDS-PAGE and Western Blotting with conditions described previously. Primary antibodies used were anti-rhodopsin antibody (1:500, Santa Cruz, sc-57432) and anti-SLC6A19 antibody (1:2000, HPA043207-100UL, Sigma-Aldrich). Anti-rhodopsin blots were incubated in goat anti-mouse HRP conjugate (1:10,000, Bio Rad, 1721011). Anti-SLC6A19 blots were incubated in donkey anti-rabbit HRP conjugate (1:10,000, A16023, ThermoFisher). Visualization was performed using a G:Box (Syngene).

Statistical analysis:

Statistical analyses were performed using GraphPad Prism 9 (version 9.1.2, GraphPad software, USA). Sigmoidal dose response curves and interpolated IC50 values were determined using Sigmoidal, 4PL model of regression analysis in GraphPad Prism. For statistical significance, the datasets were initially assessed for Gaussian distribution, based on which further analysis was performed. For datasets that followed normal distribution, unpaired t-test was used to compare two groups. Details of statistical tests used for different data sets have also been provided in figure legends.

## Supplementary Figures

**Figure S1.**
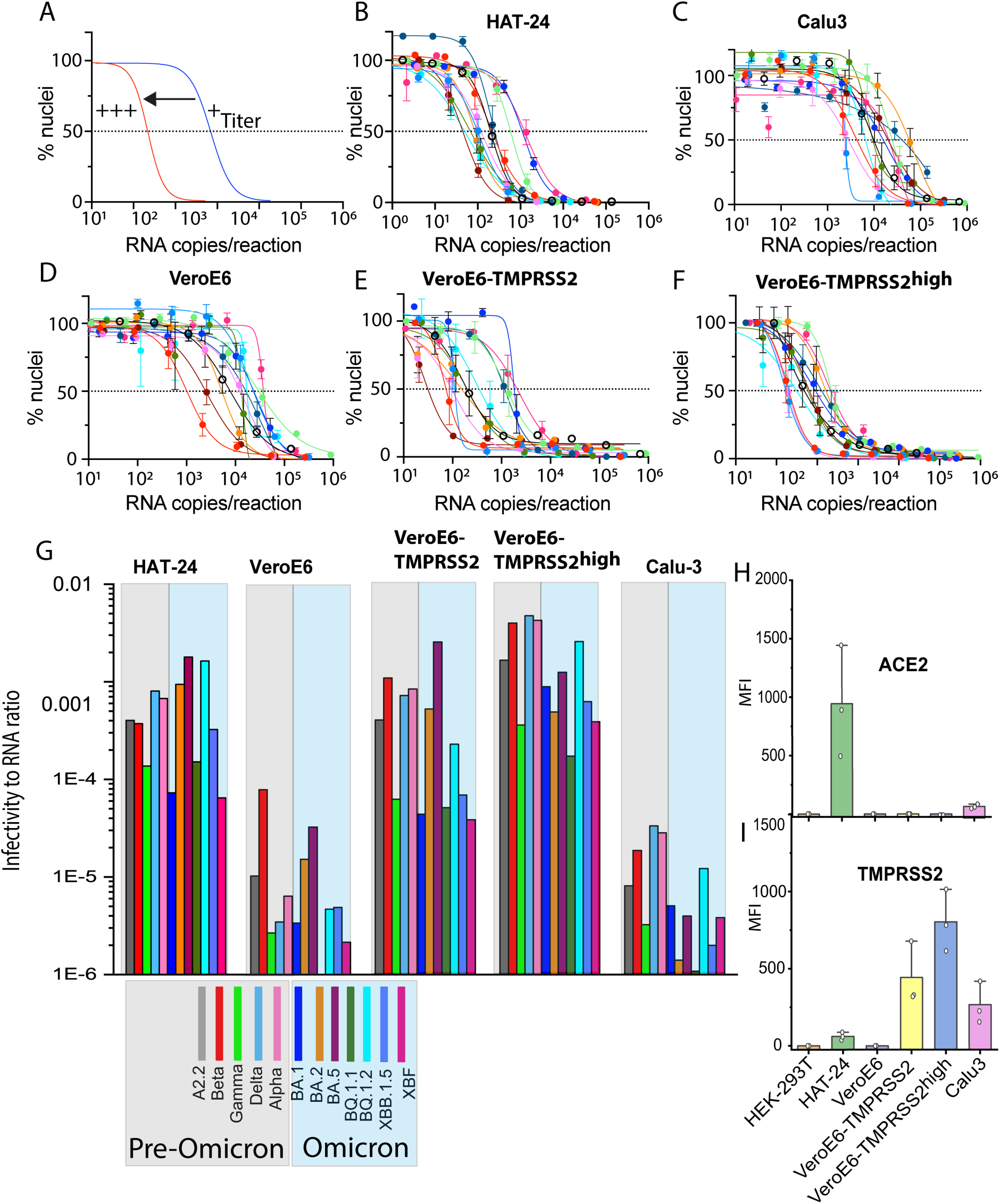
Engineered cell lines with TMPRSS2 reveal a continuum of sensitivity across all major primary SARS-CoV-2 variants A. Schematic illustrating high content datasets relative to RNA input. A shift in the sigmoidal curve to the left represents a viral isolate with a greater titer (low viral input can sustain significant cytopathic effects) which is readily enumerated by high content microscopy using live fluorescent nuclei counts. The dotted line represents 50% loss of nuclei and interpolated values can be used to determine viral titers. **B-F.** Titration curves per RNA input for the 12 primary clinical isolates highlighted in A. **B.** The Hek 239T engineered HAT-24 cell line co-expressing ACE2 and TMPRSS2, as previously described ^17^. **C.** Calu-3 human lung adenocarcinoma cell line. **D.** VeroE6 cell line. **E.** VeroE6 cell line expressing TMPRSS2 (VeroE6-TMPRSS2), as previously described^3^. **F.** VeroE6 cell line expressing TMPRSS2 (VeroE6-TMPRSS2^High^), as previously described ^28^. **G.** Infectivity is calculated by the viral dilution requited to sustain 50% of the loss of nuclei and then expressed as an infection to RNA ratio on the Y-axis (VE50). Grey and blue shading represents pre-Omicron and Omicron lineages, respectively. Infectivity to RNA ratios are derived from data generated from C-G. and is representative of three independent experiments. **H & I.** Cell surface staining and flow cytometry of cells used herein for **I.** TMPRSS2 and **J.** ACE2. Each data point represents an independent experiment.

**Figure S2.**
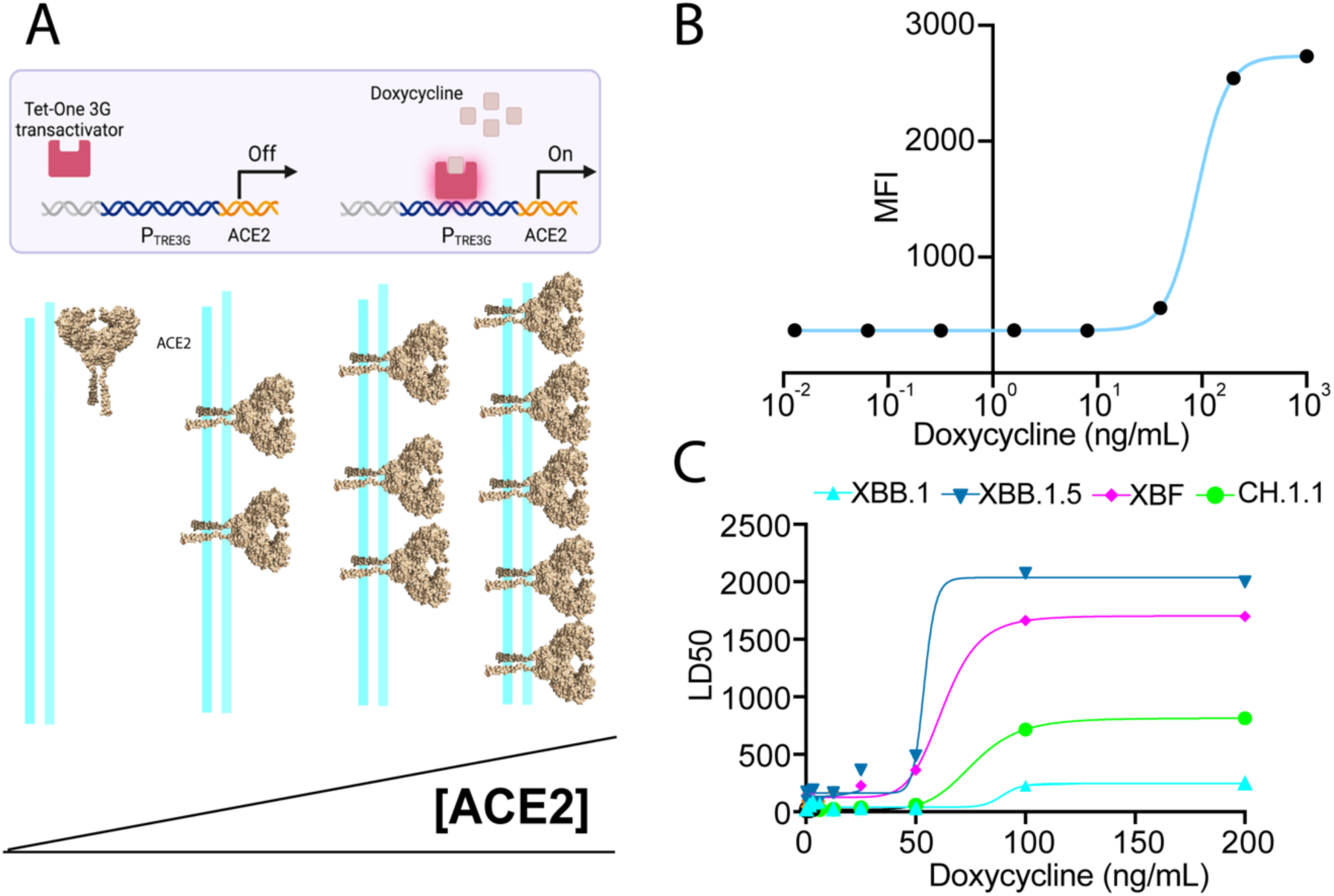
ACE2 usage by Omicron lineages using the ACE2 Affinofile assay. A. Affinofile HEK-293T cells engineered to inducibly express ACE2 using a Doxycycline sensitive promoter. B. ACE2 expression in the Affinofile system in the presence of increasing concentrations of doxycycline, as measured using surface staining for ACE2 and flow cytometry. MFI = mean fluorescence intensity. C. Viral titers for XBB.1, XBF, XBB.1.5 and CH.1.1 using the Affinofile cell line outlined in J. are generated by determining the dilution that sustains 50% cell loss in each culture (Lethal Dilution 50%; LD50).

**Figure S3.**
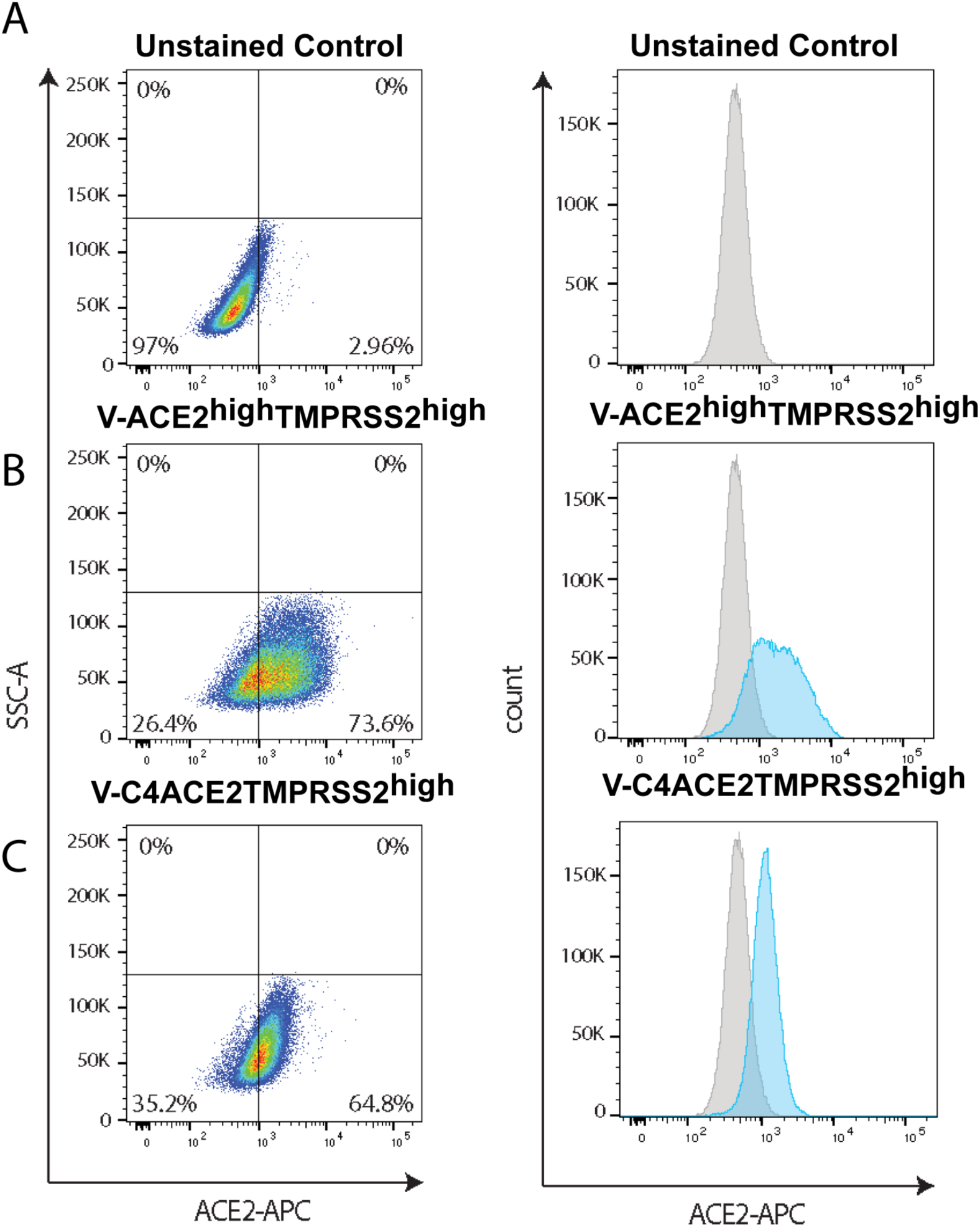
ACE2 surface expression of the VeroE6-T2 cell line with WT ACE2 and ACE2 C4 mutant (C4-ACE2). A. Unstained control. B. V-ACE2^high^-TMPRSS2^high^ stained for ACE2. C. V-C4ACE2-TMPRSS2^high^ stained for ACE2.

**Figure S4.**
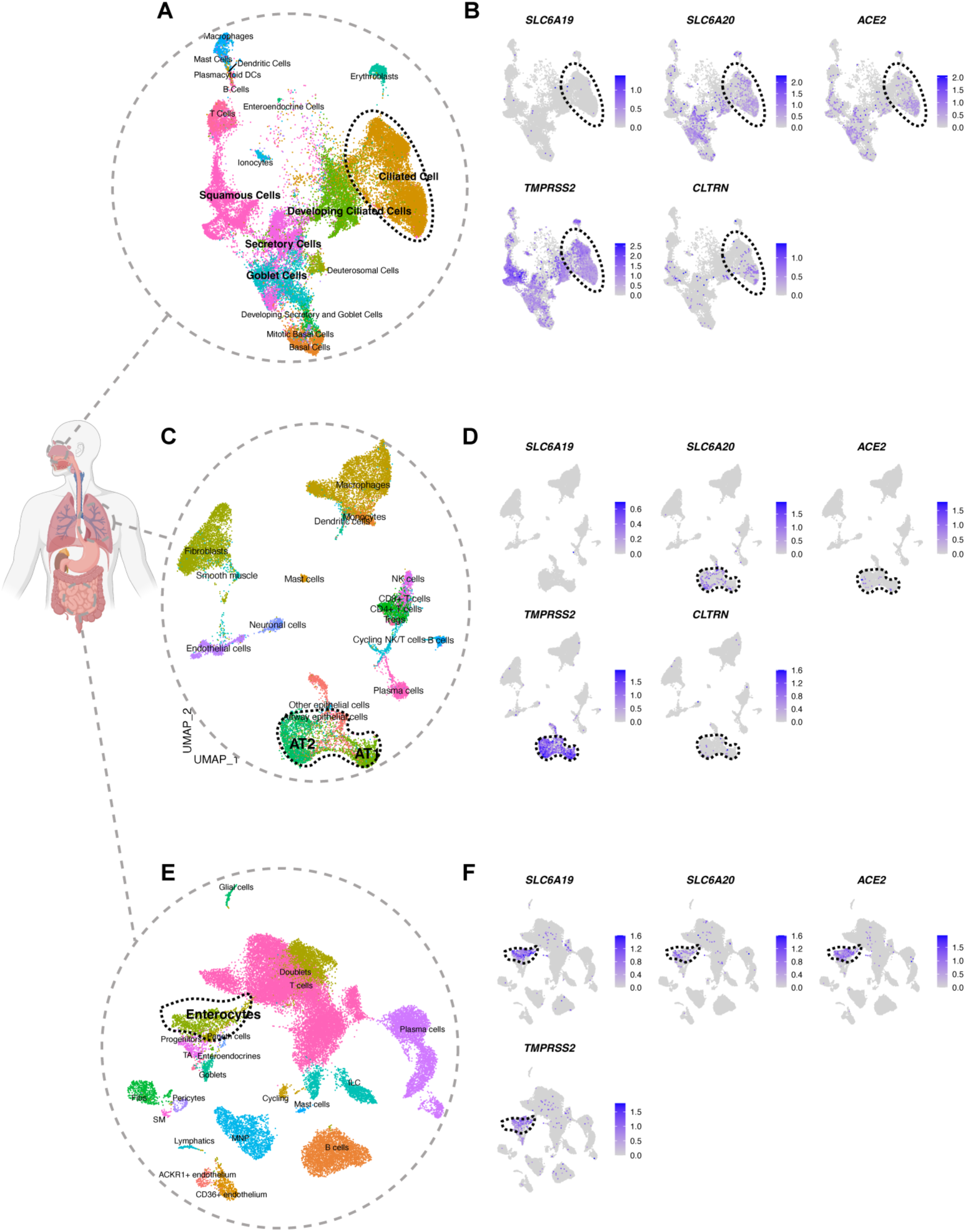
*In vivo* profiling of solute carriers SLC6A19 and SLC6A20 as additional SARS-CoV-2 entry factors alongside known entry factors ACE2 and TMPRSS2. A.-F. Uniform manifold approximation and projection (UMAP) of **A**. Nasal Epithelia. **C.** Lung and **E.** Small intestine using single nuclei RNA sequencing. Cell types are color-coded. Target cells within each tissue are highlighted in B. D. & F. in respective tissues, with expression of protein solute carriers SLC6A19 and SLC6A20 alongside the known SARS CoV-2 receptors ACE2 and TMPRSS2. Collectrin (CLTRN) is also presented as it represents and additional chaperone for SLC6A20 and can compete for ACE2 in forming a dimer of heterodimers. Of note, ACE2 single cell expression is closely aligned with SLC6A19 and SLC6A20 in the small intestine and nasal cavity respectively. In contrast, the lung is primarily defined by high levels of TMPRSS2 expression in type 2 pneumocytes.

**Figure S5.**
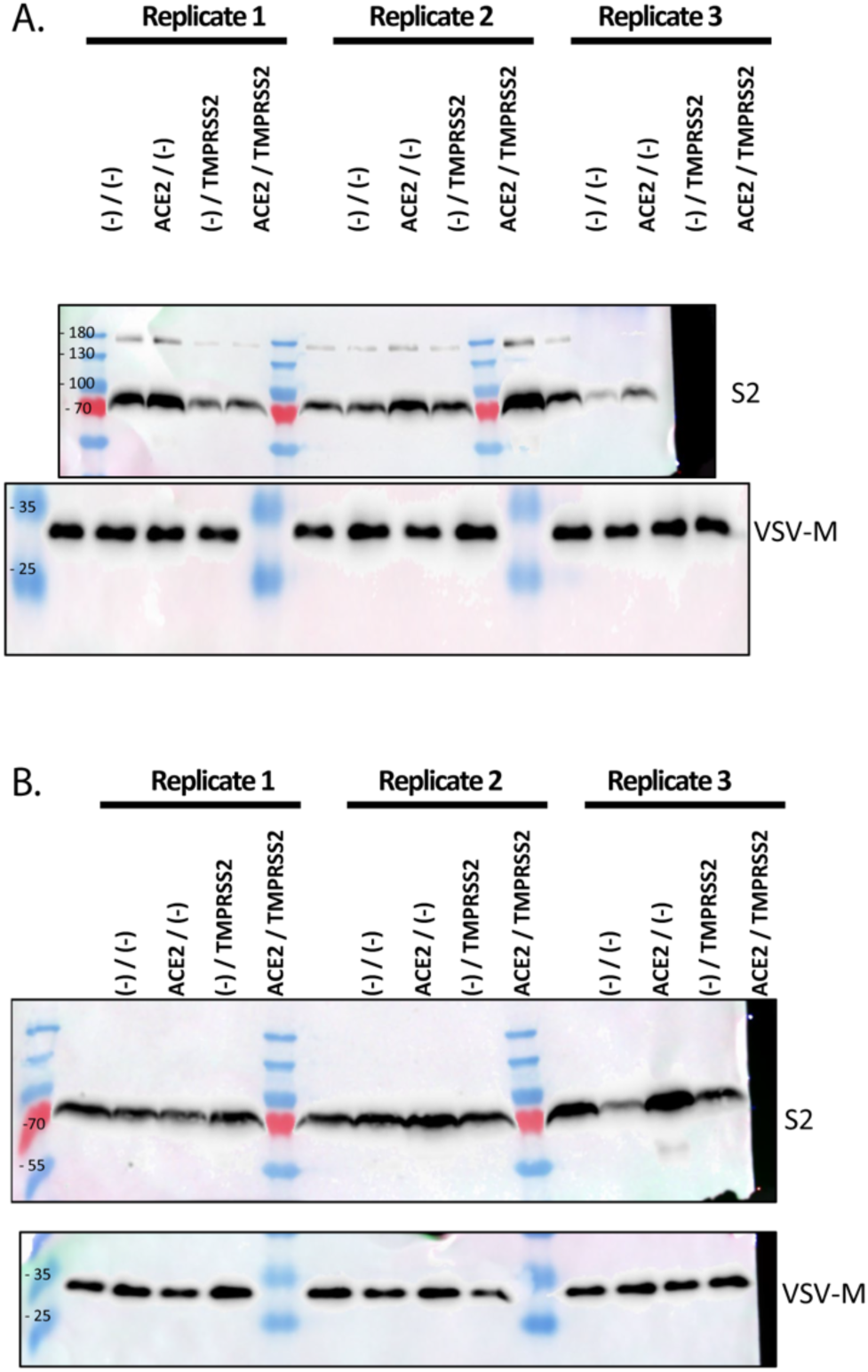
Spike S1/S2 cleavage in representative pre-Omicron and Omicron VSVg pseudotypes. **A. & B**. All receptor constructs in lentiviral constructs used herein were shuttled to the transient mammalian expression vector pcDNA6. Pseudovirus particles were then generated in the presence or absence of various receptor combinations. “(-)/(-)” indicates no receptor control “ACE2/(-)”-WT ACE2, “(-)/TMPRSS2”-TMPRSS2 and “ACE2/TMPRSS2”-combined ACE2 and TMPRSS2 expression during pseudoparticle production in HEK-293T cells. Three days post transfection, pseudoviruses were pelleted via ultracentrifugation for 90’ at 100,000xg through 20 % sucrose cushion. Pellets were then lysed and subjected to Western blotting. Here pseudoviral proteins-VSV-M represents the viral loading control and spike protein is detected for S1/S2 cleavage using the S2 antibody as described herein for primary isolates. **A**. Represents an early circulating Pre-Omicron B-Clade, whilst **B**. represents the more recent Omicron KP.3 lineage. Size of molecular weight markers are presented to the left of Western blots and are in KDa. In Both A. and B. three independent viral pseudotype preparations are presented.

**Figure S6.**
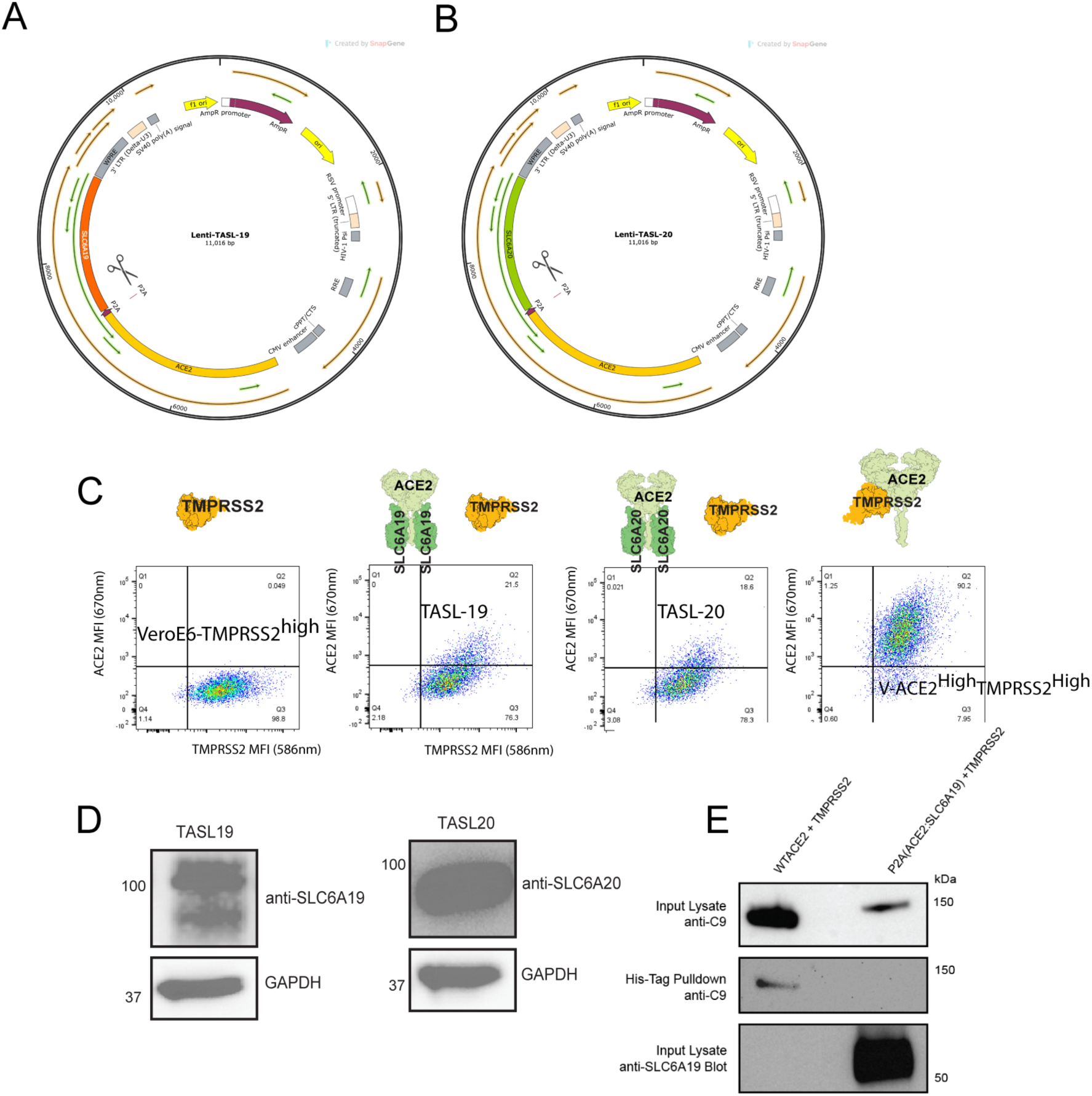
Establishment of the TMPRSS2-ACE2-Solute Carrier Cell lines TASL-19 and TASL-20. A. & B. Maps of bicistronic lentiviral constructs to enable equimolar protein expression of ACE2 and A. SLC6A19 or B. SLC6A20. Scissors highlight the P2A cleavage site that liberates ACE2 and either solute carrier. **C.** Development of the TASL-19 and TASL-20 cell lines. Here lentiviral constructs from A & B have been used to genetically modify the VeroE6-T2 cell line with either ACE2 and SLC6A19 or B. SLC6A20. To validate integration and expression, cells were stained using ACE2 and also TMPRSS2. The parental cell line VeroE6-T2 and the ACE2-VeroE6-T2 are presented as controls. In right histograms data is presented for three independent stains for ACE2 and TMPRSS2. **D.** Given the close proximity of Solute Carriers to the cell membrane, we generated cell lysates from both TASL-19 and TASL-20 clones to confirm expression of each at the protein level. Given their expression is through a bicistronic vector, ACE2 expression alone can act as a surrogate for quantitative solute carrier expression in each cell line**. E.** TMPRSS2 pull-down of ACE2 only in WT versus ACE2+ SLC6A19. Pull-down of TMPRSS2 is via His-tag pull-down and then blotting for ACE2 using the C9 tag. Experiments were carried out using transient expression in the Hek 239T cell line with identical constructs expressed in the pcDNA6 CMV vector.

